# Bacteriophages of the predominant coral symbiont *Endozoicomonas*: novel models for coral holobiont interactions

**DOI:** 10.1101/2025.10.29.685362

**Authors:** Po-Shun Chuang, Mo Chen, Pei-Wen Chiang, Hsing-Ju Chen, Feng-Chi Wang, Yu-Hsiang Chen, Sheng-Ping Yu, Fu-Chi Chen, Kai-Ning Shih, Yung-Pei Chang, Sim Lin Lim, Hsin-Feng Chang, Wann-Neng Jane, Sen-Lin Tang

## Abstract

Phages are important symbionts in corals that modulate the community and functions of other symbiotic bacteria. Although phages infecting coral pathogens have been reported, no phage targeting beneficial microorganisms in corals has been isolated to date. From seawater near *Acropora* and *Stylophora* corals, we isolated the first bacteriophages (designated EmPhiA and EmPhiS) that infect *Endozoicomonas montiporae* CL-33, a model strain of the coral-prevalent and predominant *Endozoicomonas* bacteria. Electron microscopic observations of both phages showed *Myovirus*-like morphology and head sizes characteristic of jumbophages, with cryo-electron microscopy reveals long whiskers unprecedent in known phages. Genetically, these phages shared 99.21% genome similarity and are distant from known prokaryotic viruses, suggesting that they represent a novel viral species, which we name *Encorevirus taiwanensis*, in a novel family *Encoreviridae*. The small burst sizes of these phages (13.14 PFU/cell for EmPhiA and 21.4 PFU/cell for EmPhiS) potentially enable continuous coexistence of them with host bacteria within corals, making them putative core members of coral holobionts. Furthermore, host range test showed that EmPhiA and EmPhiS infect both *Endozoicomonas* bacteria isolated from stony and soft corals, implying their presence in a broad spectrum of host marine invertebrates. Using EmPhiS, we also investigated phage-bacterium interaction during its infection of *E. montiporae* CL-33. Interestingly, in addition to modulation of host cellular machinery, we found expression of several tellurium resistance proteins by EmPhiS during infection, which may provide the host additional stress resistance. These phages provide a novel model that will greatly advance our understanding of coral-*Endozoicomonas*-phage interactions.

## Introduction

Interactions of symbiotic microorganisms determine the physiology of a coral and its ecological functions. Viruses are essential components of coral holobionts but their contribution is less studied compared to microalgae and bacteria. By infecting and lysing bacteria in coral holobionts, viruses modulate bacterial communities associated with corals [1–3]. Viruses in reef seawater may also contribute to corals by lysing bacteria and phytoplankton in the water column, making energy and nutrients available to corals [4, 5]. In addition, some viruses carry extra metabolic genes that are expressed during infection or incorporated into host genomes (lysogenic infection), by which they alter ecological functions of other coral-associated microorganisms [6, 7].

*Endozoicomonas* bacteria are widely perceived as beneficial microorganisms for corals due to their prevalence in healthy corals [8, 9]. Recent studies have begun to decipher interactions between *Endozoicomonas* bacteria and their host corals, which may involve eukaryotic-like genes and the type III secretion system and may contribute to sulfur and phosphate cycles in coral reef ecosystems [10–12]. Interestingly, studies have identified many prophage genes in *Endozoicomonas* genomes, suggesting a potential role of phages in *Endozoicomonas* eco-functions [3, 13, 14]. However, contemporary research of coral-associated viromes has focused mostly on eukaryotic viruses and pathogenic phages that show potential applications in treating coral diseases. The influences of phages on mutualistic bacteria such as *Endozoicomonas* remain unknown.

In this study, we isolated two phages from coral reef seawater that infect *Endozoicomonas montiporae* strain CL-33^T^, one of the most studied *Endozoicomonas*. With biochemical tests, high-resolution electron microscopy, and genome sequencing, we characterized their morphological, physiological, and genetic features. Using one of the isolated phages, we also examined transcriptome dynamics during its infection of *E. montiporae* CL-33. Findings from this study offer a novel model for studies of symbiosis among phages, *Endozoicomonas*, and corals.

## Methods and Materials

### Seawater collection

In August 2019, we collected seawater from the surfaces of *Stylophora pistillata* and *Acropora sp.* colonies at Kenting National Park in southern Taiwan (1 L/species) using 50 mL syringes. Collected seawater was successively filtered through 10-µm and 0.22-µm membrane filters to remove plankton and bacteria. Filtrates were concentrated to 50 mL using a 50-kDa filter with the Labscale TFF system (Merck Millipore, Germany) and preserved at 4°C.

### Purification of Endozoicomonas phages

To enrich *Endozoicomonas*-infecting phages, 40 mL of log-phase cultured *E. montiporae* CL-33 were inoculated with 10 mL of seawater filtrate and incubated overnight at 25°C with 200 rpm shaking. All bacteria in this study were cultured in MMBv4 medium, as described by Ding et al [13]. A plaque assay was performed using the standard double-layered agar method, with *E. montiporae* CL-33 as the host. After one-day incubation at 25°C, plaques were picked and incubated overnight at 4°C in 1 mL of MMBv4 to recover bacteriophages. The process was performed more than three times to obtain pure isolates of *E. montiporae* CL-33 phages (designated EmPhiA and EmPhiS for those isolated from seawater near *Acropora* and *Stylophora* corals, respectively).

### Host range, temperature, and chloroform sensitivity tests

Host ranges of isolated bacteriophages and their temperature and chloroform sensitivity were tested using spot assays. For the host range test, 10 *Endozoicomonas* strains isolated from different marine invertebrates were employed as hosts (Table 1). For the temperature sensitivity test, 1 mL of bacteriophage (10¹LJ PFU/mL) was pre-incubated at 31°C for 2 h before subjected to a spot assay, with a control group comprising non-stress phages.

**Table 1.** Host ranges of EmPhiA (PhiA) and EmPhiS (PhiS) against 10 *Endozoicomonas* strains. Sources and references of bacterial isolates are listed.

| Host | PhiA | PhiS | Source | Reference |
| --- | --- | --- | --- | --- |
| *E. elysicola* MKT110 | - | - | Sea slug | Kurahashi and Yokota [35] |
| *E. montiporae* CL-33<sup>T</sup> | + | + | Stony coral | Yang et al [36] |
| *E. euniceicola* EF212<sup>T</sup> | + | + | Soft coral | Pike et al [37] |
| *E. gorgoniicola* PS125<sup>T</sup> | - | - | Soft coral | Pike et al [37] |
| *E. numazuensis* HC50 | - | - | Sponge | Nishijima et al [38] |
| *E. atrinae* WP70 | - | - | Molluscs | Hyun et al [39] |
| *E. acroporae* Acr-14<sup>T</sup> | - | - | Stony coral | Sheu et al [40] |
| *E. coralli* Acr-12<sup>T</sup> | - | - | Stony coral | Chen et al [41] |
| *E. ruthgatesiae* 8E | - | - | Stony coral | Chiou et al [10] |
| *E. penghunesis* 4G | - | - | Stony coral | Tandon et al [42] |

Chloroform sensitivity was tested by mixing 500 µL of bacteriophage solution (10LJ PFU/mL) with 0 µL (control group), 5 µL (1%), 50 µL (10%), or 500 µL (50%) of chloroform. Mixtures were incubated at 25°C for 30 min and bacteriophage-containing supernatants were collected for a spot assay by centrifuging at 3,000 xg for 10 min at 4°C. Phage infectivity was determined by plaque formation in the spot assay after 2 days of incubation at 25°C except for the temperature sensitivity test, which was also incubated at 31°C.

### One-step growth test

A one-step growth experiment was conducted by inoculating *E. montiporae* CL-33 with bacteriophages at a multiplicity of infection (MOI) of 0.1. Phage absorption was allowed for 1 min, followed by centrifuging at 12,000 xg for 30 sec at room temperature. Supernatant (unabsorbed phage particles) was subjected to a spot assay to estimate phage absorption rates, while phage-infected bacterial cells were resuspended in 100 mL of MMBv4 medium and incubated at 25°C with 200-rpm shaking. The phage-bacteria mixture (1 mL) was sampled at 30-min intervals for 150 min, filtered through a 0.22-µm membrane, and subjected to a spot assay (at 25°C) to estimate phage titer dynamics. Both temperature and chloroform sensitivity tests and the one-step growth experiment were performed in triplicate.

### Transmission electron microscopy (TEM) and cryogenic electron microscopy (cryo-EM)

TEM samples were prepared by applying 5 µL of isolated bacteriophages onto a glow-discharged grid and were negatively stained with 2% uranyl acetate. TEM observation was performed on a FEI Tecnai G2 Spirit TEM (Thermo Fisher, USA) and images were taken with a 4K x 4K Gatan Orius CCD camera (Gatan, USA) with a defocus range of -1.5 to -2.5 µm. For cryo-EM samples, 4 µL of isolated bacteriophages were applied to a glow-discharged 200-mesh Quantifoil R2/1 holey carbon grid, which was then blotted at 100% humidity at 4°C for 3.5 sec and plunge-frozen in liquid ethane using a FEI Vitrobot Mark IV System (Thermo Fisher, USA). Cryo-EM observation was performed on an FEI Tecnai F20 TEM (FEI company, USA) and Talos Arctica cryo-TEM (Thermo Fisher, USA). Cryo-EM images were taken with a Falcon LJ detector with a defocus range of -0.5 to -2.5 µm (Thermo Fisher, USA) and data were processed using cisTEM. TEM and cryo-EM sample preparation and data processing were conducted by the Electron Microscope Division, Cell Biology Core Lab at the Institute of Plant and Microbial Biology, Academia Sinica, Taiwan (herein referred EM Division).

### DNA extraction and whole genome sequencing

Bacteriophage isolates were first treated with DNase I to remove free DNA from host bacteria. Viral particle purification was carried out using a CsCl step gradient (density: 1.35-1.5 g/mL) with a Beckman Optima L-80 XP ultracentrifuge (Beckman Coulter Inc., USA) at 4°C and 87,000 xg for 2 h, after which the phage particle-containing band was extracted with a needle and stored at 4°C. Viral DNA was extracted using the formamide/cetyltrimethylammonium bromide method [15] and was subjected to sequencing on Illumina HiSeq 2500 and Oxford Nanopore sequencing systems (Yourgene Bioscience, Taiwan).

### Genome assembly

Genome assembly of isolated bacteriophages was performed with long reads from Oxford Nanopore sequencing using the software Flye v2.8 [16]. Short reads from Illumina HiSeq 2500 were trimmed and filtered using trimmomatic v0.39 [17] with the following parameters: ILLUMINACLIP:TruSeq3-PE-2.fa:2:30:10:3: TRUE LEADING:10 TRAILING:10 SLIDINGWINDOW:5:15 MINLEN:50 and mapped to the assembled genomes using BWA-MEM [18]. Aligned reads were employed to further polish the assembled genomes using Pilon v1.23 [19].

### Genome annotation

Open reading frames (ORFs) were predicted using VIBRANT v.1.2.1 [20]. Annotation was conducted using both VIBRANT v1.2.1 and BLASTp against the non-redundant protein sequences (nr), RefSeq Select proteins (refseq_select), and megagenomic proteins (env_nr) databases in NCBI, with criteria set as E-value ≤10^-5^, alignment coverage ≥50%, identity ≥25%, and bit score ≥150. GC skew was estimated using GenSkew (Webskew: https://genskew.csb.univie.ac.at/) with default settings. Transfer RNA (tRNA) prediction was performed using tRNAScan-SE v2.0.7 [21] with the general tRNA model and fast scan mode. Repeated genetic elements were searched using the *Find Repeat* function in Geneious 10.2.6 [22] with a threshold of minimum repeat length = 20 bp and maximum mismatch = 0%. Genome annotation, GC-cumulative skew curves, predicted tRNA, and repeated elements were incorporated into genome assemblies using Geneious 10.2.6.

### Phylogenetic and comparative analysis

Genome similarity between isolated bacteriophages and known viruses in the NCBI database (taxid:10239) was estimated using BLASTN with an E-value <1e^-5^ and coverage >50%. A genome-based, proteomic phylogenetic tree was reconstructed with isolated bacteriophages and 5632 reference genomes of prokaryotic dsDNA viruses using ViPtree v4.0 [23]. A simplified phyogenetic tree was also reconstructed with 316 randomly selected genomes from the reference pool. The three reference genomes most similar to isolated bacteriophages (*Vibrio* phage eugene 12A10, *Klebsiella* phage vB_KpnM_KB57, and *Proteus* phage Mydo) and 3 random genomes in the branch neighboring isolated bacteriophages (family *Demerecviridae*; *Aeromonas* phage AhSzw-1, *Vibrio* phage vB_VpsPG07, and *Vibrio* phage Pontus) were selected to generate alignment plots against our bacteriophages using ViPtree v4.0. Proteomic trees for individual marker genes were constructed for DNA polymerase, terminase, helicase, tail fiber, and RNase H. Using EmPhiA genes as queries in BLASTP searches, the top 10 hits for each marker gene in the NCBI database were selected as references for gene-based phylogenetic analysis. Marker genes from phages in the viral families *Ackermannviridae*, *Chaseviridae*, *Demerecviridae*, and *Pseudomonas* phage PAK_P1 (Genus: *Pakpunavirus*, Family: unassigned) were also included in the analysis. Homologs of EmPhiA and EmPhiS genes in closely related viral families (*Ackermannviridae*, *Chaseviridae*, *Naomviridae*, and *Demerecviridae*) were searched using Coregenes 5.0 [24] with the criterion of an E-value <1e^-5^.

### Transcriptome dynamics during EmPhiS infection

To synchronize phage infection, EmPhiS was mixed with *E. montiporae* CL-33 at a MOI of 3. Phage absorption was allowed for 10 min, after which free phage particles were isolated by centrifuging at 13,200 rpm for 2 min to estimate the absorption rate. Infected bacteria were resuspended in 100 mL MMBv4 medium and incubated at 25°C with 200 rpm shaking. A non-phage control group was prepared by inoculating the same volume of MMBv4 medium. The experiment was performed with 5 replicates, from which 10 mL of phage-bacterial culture were sampled at an interval of 30 min for 2 h. Collected samples were centrifuged at 3,000 rpm for 15 min. Bacterial pellets were subjected to RNA extraction following the TRIzol method and to TEM after cryo-fixation, while supernatants were employed to a plaque assay for phage titer estimation. Extracted RNA was submitted to the NGS High Throughput Genomics Core in Academia Sinica, Taiwan for library preparation (Illumina Stranded Total RNA Prep kit with Ribo-Zero Plus) and sequencing (Illumina HiSeq 2500, single-end 150 bp). Cryo-fixation and TEM were conducted by the EM Division, with detailed procedure provided in Supplementary Material 1.

### Transcriptomic data analysis

The quality of RNA sequencing data was first checked using FastQC v0.12.1 [25]. Trimmomatic v0.39 was employed to remove sequencing adaptors. Barrnap v0.9 [26] was used to search for remaining ribosomal RNA reads in the dataset, which were format-converted using Bedtools v2.31.1 [27] and removed by Hisat2 v2.2.1 [28]. Following decontamination, bacterial and phage reads were assigned by matching against corresponding genome assemblies using Hisat2 v2.2.1. Cross-matching was also conducted to examine genome overlap between the two components. Decontaminated data were format-converted using SAMtools v1.20 [29] and gene expression was determined using the *featureCounts* function in Subread v2.0.6 [30]. Differential gene expression was estimated using DESeq2 v1.44.0 [31] with the apeglm algorithm for data shrinkage [32]. Criteria for significant differences were set as: average read counts >10, |log fold change| >1, and an adjusted *p*-value <0.05. Kyoto Encyclopedia of Genes and Genomes (KEGG) enrichment analysis was performed with eggNOG mapper 5.0 v2.1.1.2 [33] and clusterProfiler v4.12.6 [34] with criteria of *p* <0.05 and q-value <0.2.

## Results

### Isolation and characterization of Endozoicomonas phages

From seawater near *Acropora* sp. and *S. pistillata*, we successfully isolated two bacteriophages, designated EmPhiA and EmPhiS, respectively (Fig. 1a,b,f,g). Under TEM observation, both phages showed an icosahedral capsid, collar, and a long contractile tail with tail fibers (Fig. 1c,d,h,i). Cryo-EM further revealed whiskers 110-130 nm long around the collar in both phages (Fig. 1e,f,j,l). Based on cryo-EM images, the head capsid of EmPhiA was 131.2±1.6 × 122.4±2.0 nm and the tail was 259.2±1.6 × 28.4±0.8 nm, while for EmPhiS the head was 133.7±2.0 × 121.1±1.8 nm and the tail was 260.6±2.6 × 28.0±1.0 nm. Both phages infect *E. euniceicola* EF212 in addition to the original host, *E. montiporae* CL-33 (Table 1). Compared to EmPhiS, EmPhiA showed higher sensitivity to thermal stress and chloroform (Fig. 2a,b). A one-step growth test also revealed a difference between the two phages (Fig. 2c,d), with EmPhiA showing a latent period of 60 min and a burst size of 13.14 PFU/cell in an infection cycle of approximately 90 min (absorption rate = 47.1%), whereas a latent period of 30 min and a burst size of 21.4 PFU/cell were found in EmPhiS (infection cycle = 90 min; absorption rate = 55.3%).

**Figure 1.**
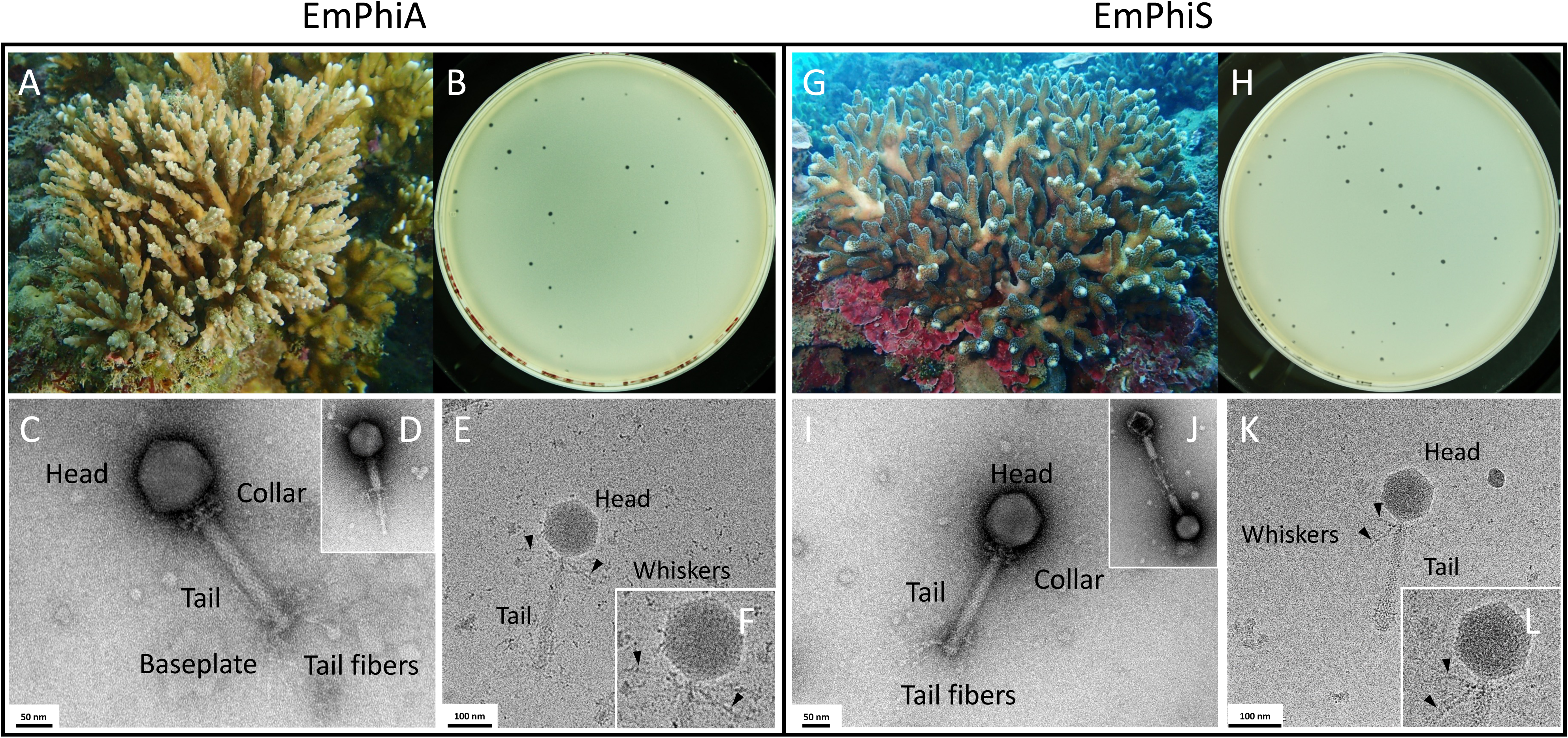
Isolation and morphology of phages EmPhiA and EmPhiS from seawater near *Acropora* sp. (a) and *Stylophora pistillata* (g) colonies, respectively. Both phages form small, round plaques on *Endozoicomonas montiporae* CL-33-base plates (b,h). TEM images reveal head-and-tail morphology with a contractile tail (insets) in both EmPhiA (c,d) and EmPhiS (i,j). Cyro-EM images reveal long whiskers (black arrows) around the collar in EmPhiA (e,f) and EmPhiS (k,l).

**Figure 2.**
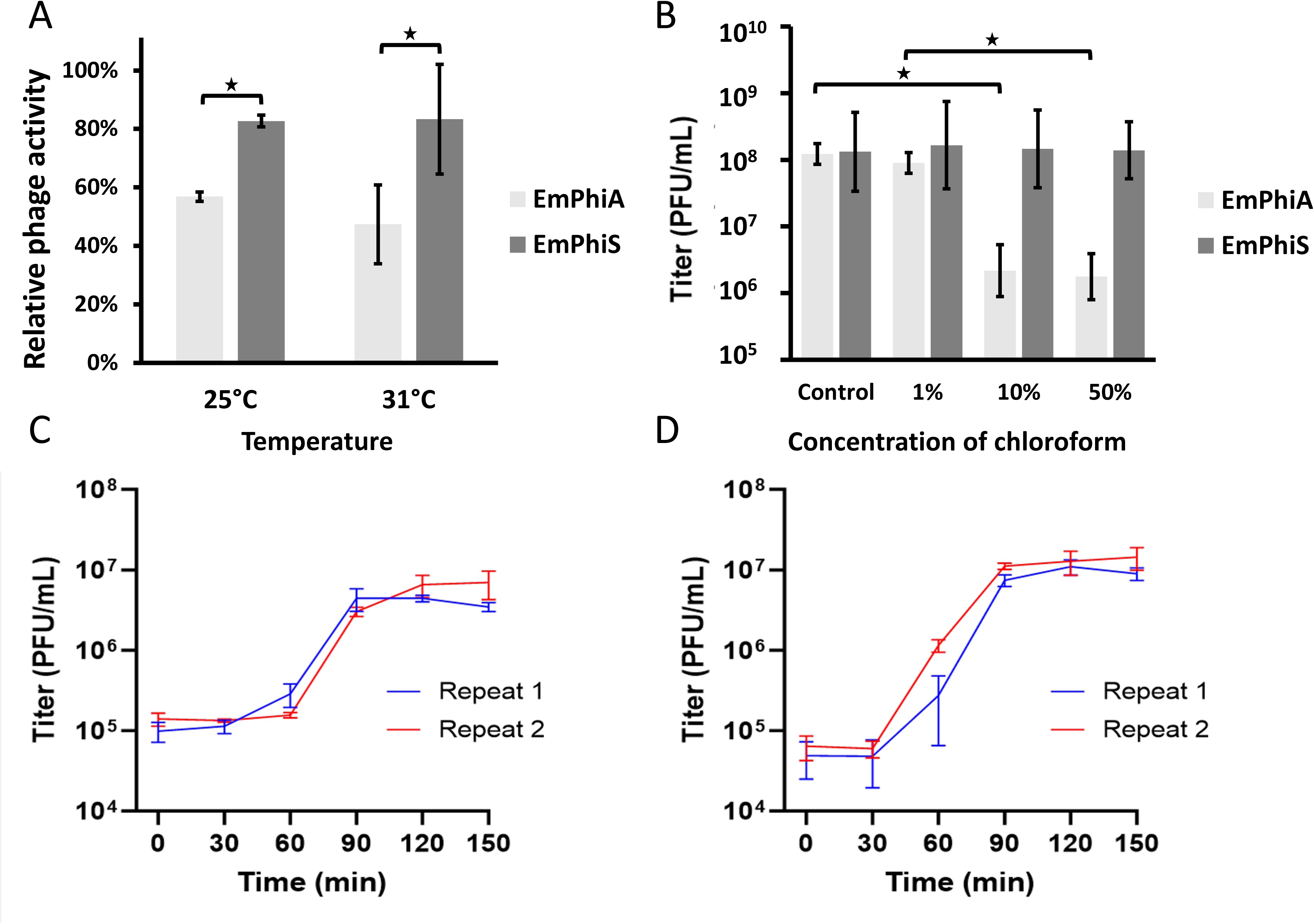
Biochemical characterization of isolated bacteriophages. (a) Sensitivity to thermal stress. (b) Sensitivity to chloroform. (c) One-step growth of EmPhiA. (d) One-step growth of EmPhiS.

### Bacteriophage genome assemblies and annotation

Using both Illumina HiSeq (EmPhiA: 0.62 Gb; EmPhiS: 0.52 Gb) and Oxford Nanopore (EmPhiA: 21.93 Gb; EmPhiS: 22.22 Gb) sequencing data, we assembled a single contig for each isolated bacteriophage (EmPhiA: 185,747 bp, EmPhiS: 186,253 bp). Both genomes showed 47.4% GC content and possessed 255 predicted ORFs, with 119 EmPhiA and 120 EmPhiS ORFs matching known or hypothetical proteins in databases. Functionally, ORFs that matched known proteins can be roughly categorized into five groups: 1) infection and lysis, e.g., cell wall hydrolase, antirepressors, and cAMP PDE, 2) nucleotide metabolism and replication, e.g., PolA, Rpo, and primase, 3) transcription regulation, e.g., tellurium resistance proteins, Rha, and Rho, 4) energy metabolism, e.g., Nampt, UroD, and MoeB, and 5) structure assembly and morphogenesis, e.g., terminase large subunit, major capsid protein, and tail fiber protein.

EmPhiA and EmPhiS showed 99.21% genome similarity and shared similar genomic organization (Fig. 3). Accumulated GC skew showed similar maximal (EmPhiA: 49,210 bp, EmPhiS: 49,290 bp) and minimal (EmPhiA: 127,095 bp, EmPhiS: 127,596 bp) positions in the two phages. Most predicted ORFs shared >90% sequence identity between the phages, except for four ORFs encoding a tail fiber (85.56% identity), a hypothetical protein (unique to EmPhiA), a phage regulatory protein (only in EmPhiS), and a HNH endonuclease (unique to EmPhiS; Supplementary Table 1). Neither phage showed preferences for or depletion of specific nucleotides or oligos. In total, 24 and 25 tRNA genes were identified in EmPhiA and EmPhiS genomes, respectively, encoding all standard amino acids except for valine (missing in both phages) and isoleucine (missing in EmPhiA). Compared to tRNA genes in *E. montiporae* CL-33, EmPhiA and EmPhiS showed higher tRNA gene ratios for threonine, cysteine, alanine, tyrosine, serine, histidine, phenylalanine, tryptophan, and asparagine, and employed some anticodons less common in the host bacterium (Fig. S1; Supplementary Table 2). Over 30 repeated genetic elements were found in both phages (EmPhiA: 30; EmPhiS: 36), including 3 shared palindrome repeats of 20 bp or 22 bp and long repeats of >100 bp (one in EmPhiA and three in EmPhiS; Fig. S2).

**Figure 3.**
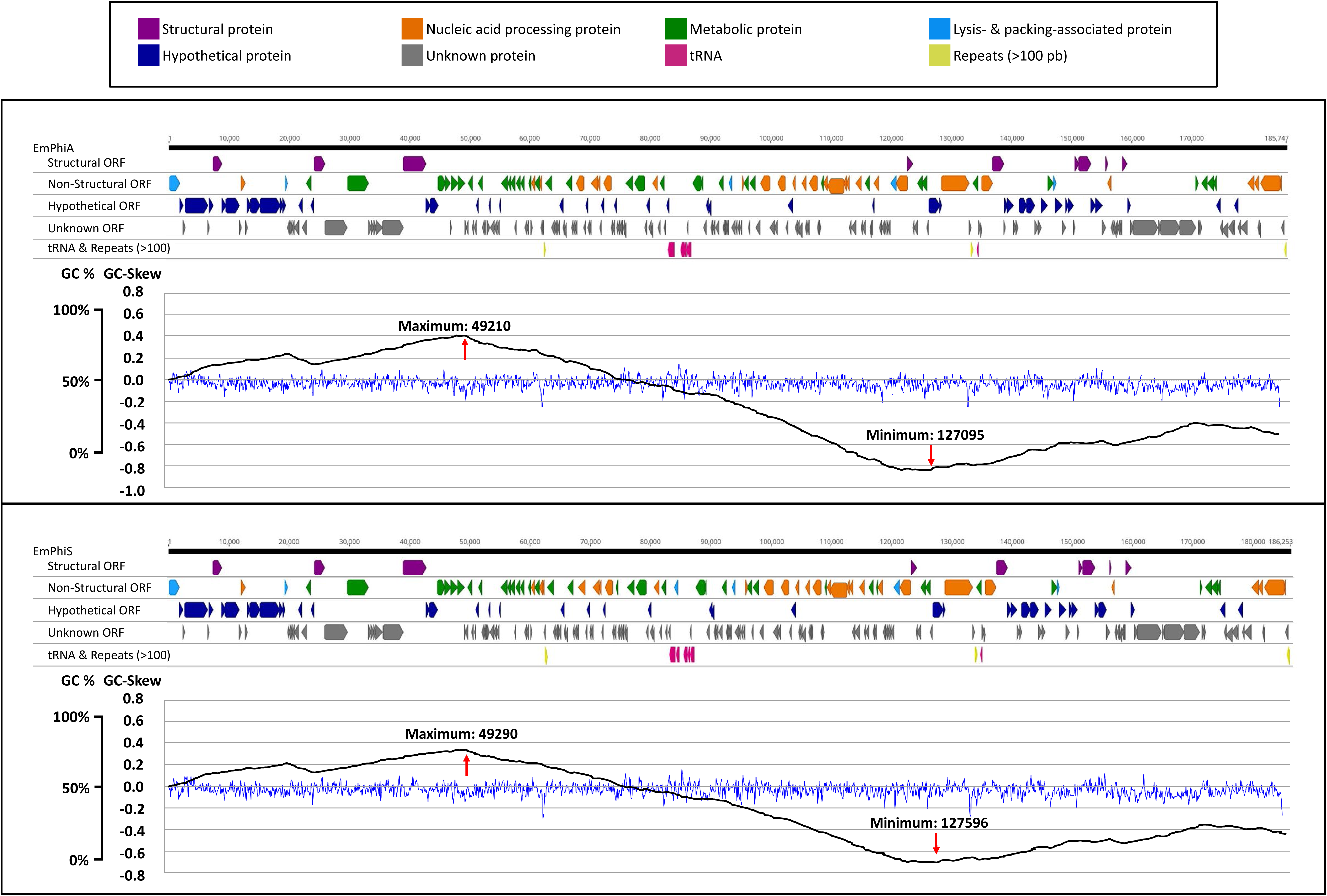
Genome maps of EmPhiA (top) and EmPhiS (bottom).

Phylogenetic analyses based on all ORFs showed monophyly of the two phages, which diverged at an early position, neighboring the viral family *Demerecviridae* with an S_G_ value <0.005 (Fig. 4). Monophyly of EmPhiA and EmPhiS was also found in marker gene-based phylogenetic trees, despite variations in its position (Fig. S3). Among closely related viral families, only *Demerecviridae* showed homologs of EmPhiA and EmPhiS genes (DNA polymerase and RNase H), whereas no homolog was found in the families *Ackermannviridae*, *Chaseviridae*, or *Naomviridae*. When aligned with the three most similar viral genomes (Fig. S4) and those of the *Demerecviridae* (Fig. S5), EmPhiA and EmPhiS showed great dissimilarity to reference sequences in terms of both gene similarity and genomic organization.

**Figure 4.**
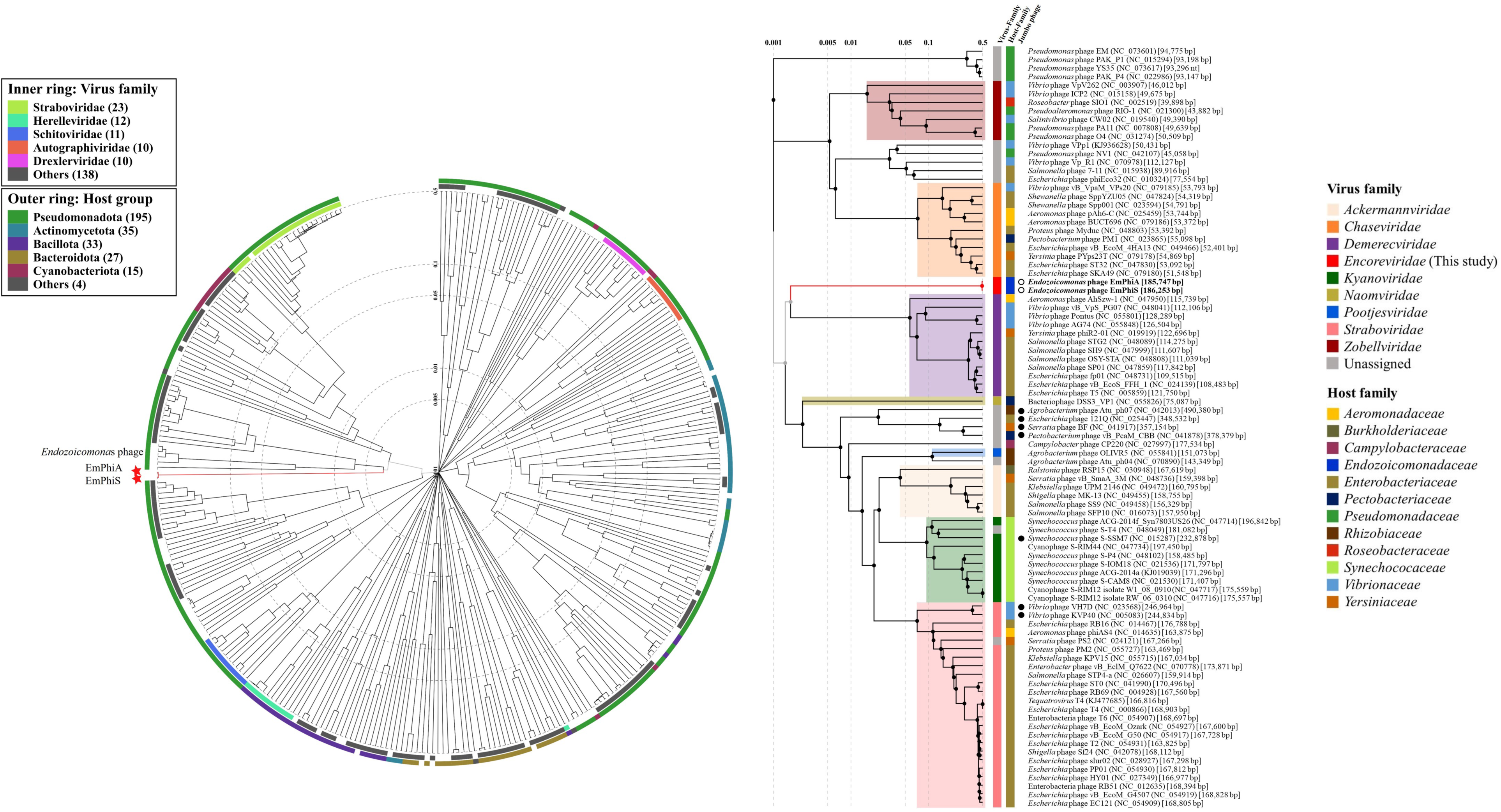
A simplified genome-based proteomic phylogenetic tree with 316 reference dsRNA prokaryotic viruses (left). EmPhiA and EmPhiS form monophyly with early divergence. A subtree for branches close to EmPhiA and EmPhiS (right). EmPhiA and EmPhiS are neighbors of the *Demerecviridae*.

### Phage-host interactions during EmPhiS infection

At an MOI of 3, one-step growth showed the same pattern as an MOI of 0.1, with an absorption rate of 26.96% and a burst size of 24.71 PFU/cell (Fig. S6). In addition to free phage particles in the culture supernatant, in our infection experiment we observed phage particles and unassembled phage structure in infected host bacterial cells (Fig. S7). From a total of 24 samples, we obtained ∼57 Gb sequences. No genome overlap was detected between the phage and host bacterium. Phage reads were merely detectable in the non-phage control but increased during infection (Fig. S8). Host transcriptomes showed clear clustering of replicates and prominent differentiation between treatments and the non-phage control from the beginning of the experiment, with the first two principal components (PC) in PCA explaining approximately 60% of the variation (Fig. 5a). On the other hand, over 90% of the variation in phage transcriptomes was explained by a single PC (PC1), which discriminated samples of 0 min from others (Fig. 5b). Clustering of replicates in the latter group, however, was less consistent. During infection, host genetic response decreased in terms of magnitude and DEG number (Fig. 6a). Due to the absence of a proper control group, phage transcriptomes were analyzed based on the expression pattern without statistical examination. Four temporal classes were identified that reflected different infection stages: early (0 min), middle (30 min), late (60-90 min) stages, and uncategorized (Fig. 6b). Because the last sampling (120 min) in our infection experiment represented the transition between the late infection stage and the onset of the next infection cycle, and because no genes were expressed specifically at this time point, it was not included in the subsequent functional analysis.

**Figure 5.**
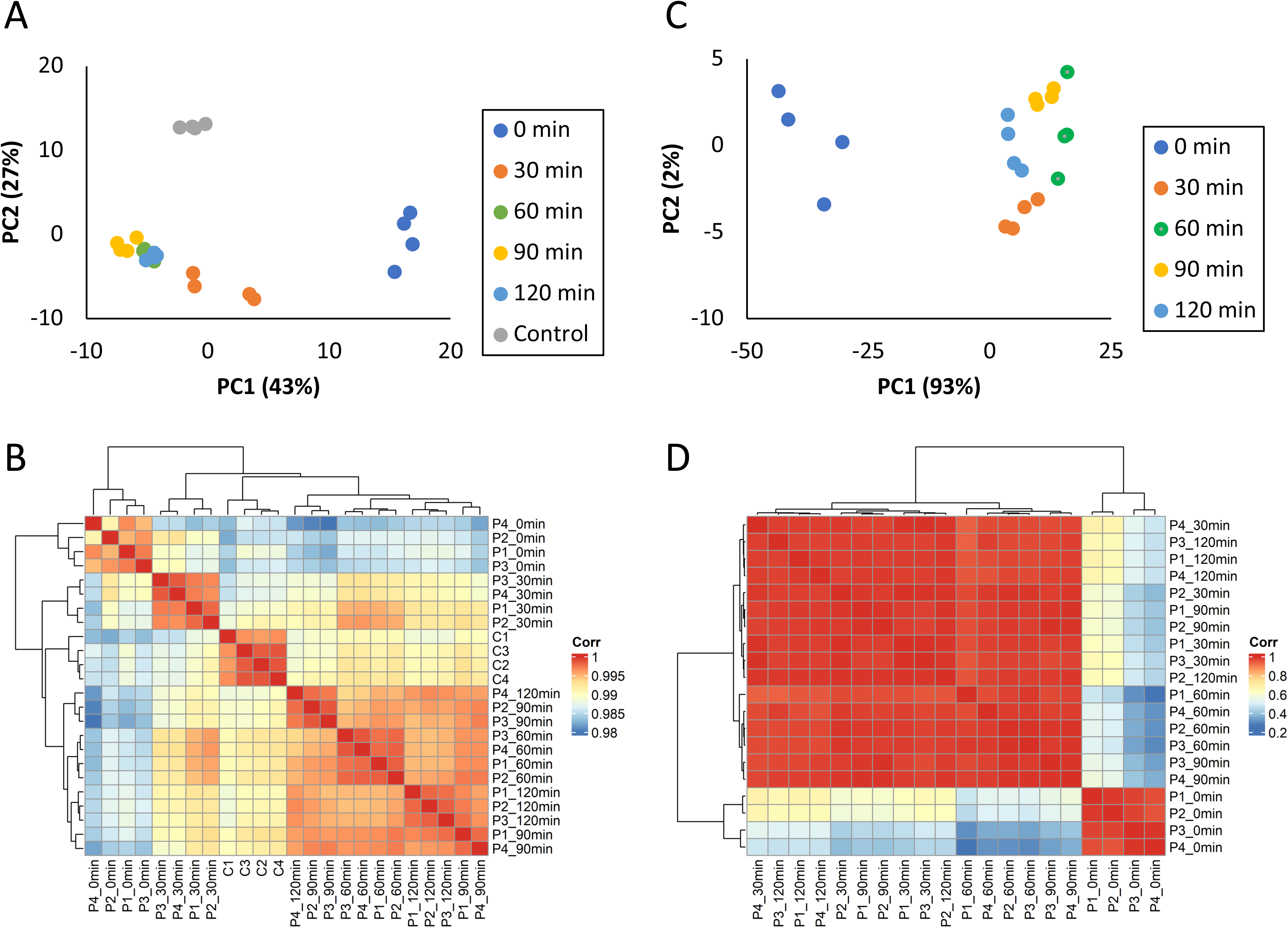
Host and phage transcriptomic dynamics during the infection experiment. (a) PCA of host transcriptome. (b) PCA of phage transcriptome. (c) Heatmap of host transcriptome. (d) Heatmap of phage transcriptome.

**Figure 6.**
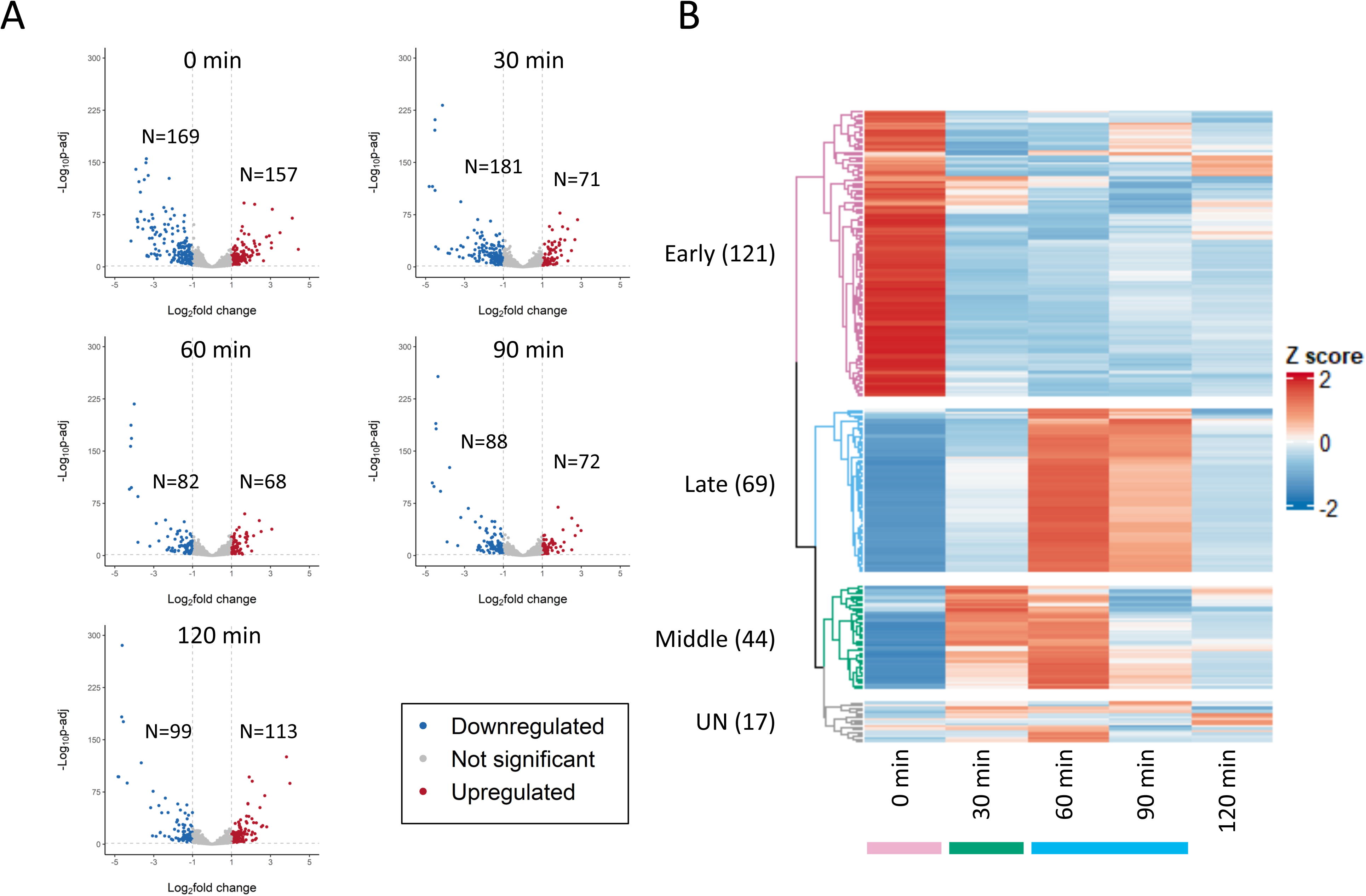
Host and phage transcriptomes during the infection experiment. (a) Volcano plots of the host transcriptome versus that of the non-phage control. (b) Phage gene expression dynamics during the infection experiment.

### Functional analysis by infection stages Early infection stage (0 min)

Due to the scarcity of phage genes with annotation, functional analysis of phage transcriptomes was performed at the gene level, disregarding statistical significance, whereas host transcriptomes were analyzed for KEGG enrichment among DEGs at specific time points. The early infection stage was characterized by expression of the highest number of phage genes, mostly related to: 1) nucleotide metabolism and replication, 2) phage infection and cell lysis, and 3) energy metabolism (Supplementary Table 3). For host DEGs (0 min vs. non-phage control), KEGG pathways like glycolysis/gluconeogenesis, glutathione metabolism, cysteine and methionine metabolism, and phosphotransferase system (PTS) were enriched among upregulated genes, whereas downregulated DEGs were enriched in pathways such as glycolysis/gluconeogenesis, TCA cycle, and flagellar assembly (Fig. 7).

**Figure 7.**
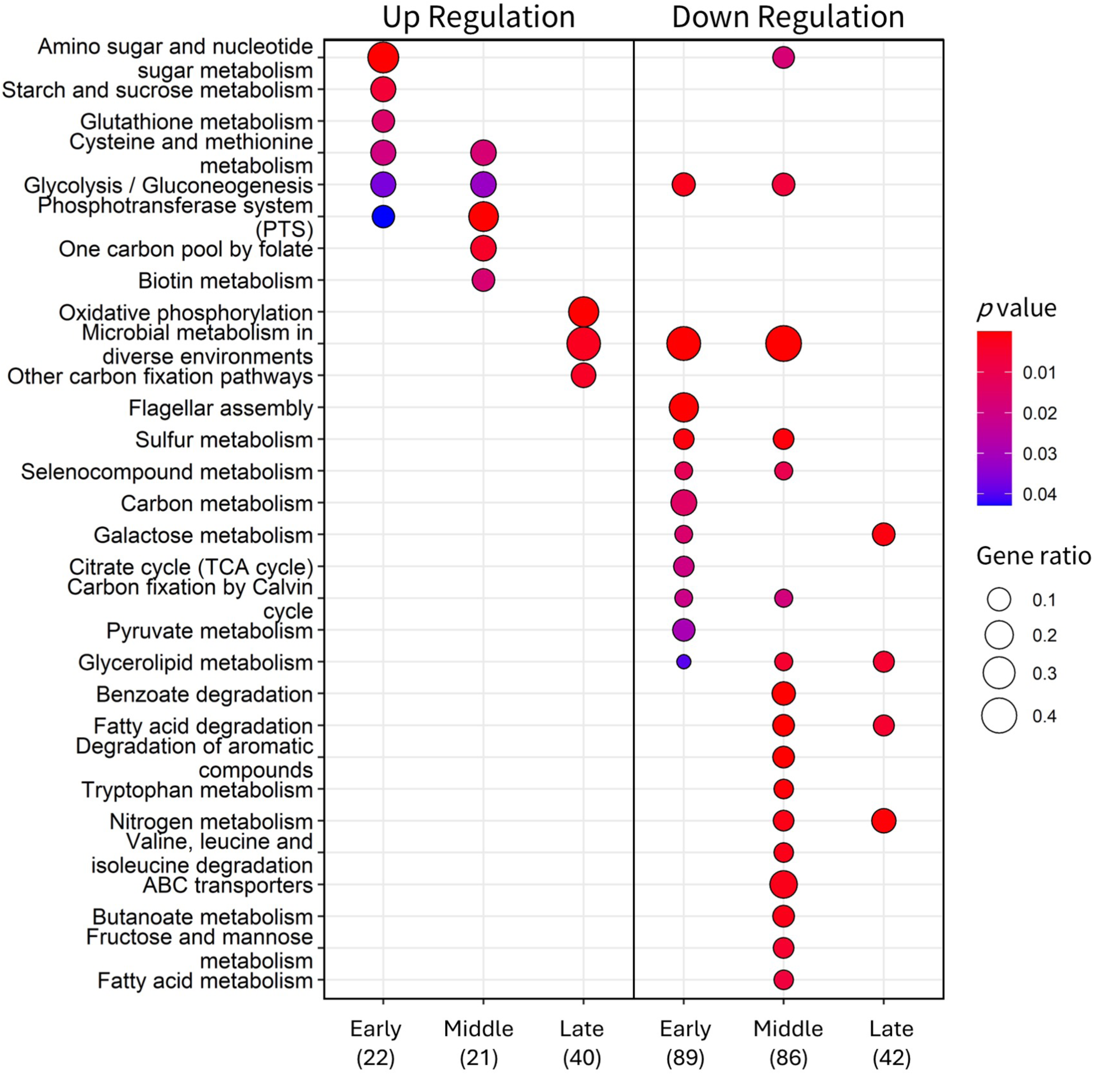
KEGG enrichment analysis of host DEGs during the infection experiment.

### Middle infection stage (30 min)

Middle-stage phage transcriptomes comprised a primase gene (DnaG), a DNA helicase, several DNA-binding protein-encoding genes, and a few other genes related to nucleotide metabolism, energy metabolism, or phage structure (Supplementary Table 4). For upregulated host DEGs at this stage (30 min vs. non-phage control), KEGG enrichment analysis showed continuous enrichment of cysteine and methionine metabolism, glyclysis/gluconeogenesis, and PTS, together with pathways such as biotin metabolism. For downregulated host DEGs, enrichment was observed in pathways related to metabolisms of nitrogen, fatty acids, and amino acids such as tryptophan, valine, leucine, and isoleucine (Fig. 7).

### Late infection stage (60-90 min)

In the late infection stage, phage transcriptomes included two RNA polymerases, an RNA polymerase sigma-G factor, two transcription factors, several genes related to nucleotide metabolism and lysis, as well as most phage morphogenesis genes (Supplementary Table 5). In host DEGs (60 min + 90 min vs. non-phage control), most KEGG pathways enriched in the early and middle stages were not identified. Instead, upregulated genes showed enrichment in oxidative phosphorylation, while downregulated genes showed enrichment in fatty acid degradation and nitrogen metabolism (Fig. 7).

## Discussion

Viruses affect coral physiology by infecting coral animals or symbiotic microorganisms and may act as part of coral immune systems by regulating associated bacterial communities [1, 2, 43]. *Endozoicomonas* is an important member of coral microbiomes that participates in carbon, sulfur, and phosphorus cycles in coral holobionts and has been proposed as an indicator of coral physiology [8, 10, 12]. In this study, we used *E. montiporae* CL-33 as a host and isolated two lytic bacteriophages, named EmPhiA and EmPhiS, from seawater collected near two common stony coral taxa, *Acropora* sp. and *Stylophora pistillata*. These are the first phage isolates that infect a putative beneficial coral symbiont. Through comprehensive morphological, biochemical, and genomic characterization, we discovered two jumbophages with unprecedent structure that represent a new viral species, which we name *Encorevirus taiwanensis*, in a new family (*Encoreviridae*). Furthermore, by examining transcriptomic dynamics during EmPhiS’s infection cycle, we offer insights into how this phage manipulates genetic machinery of the bacterial host and how this infection may affect ecology of the coral holobiont. These phages constitute a potential model to study the interaction network of coral-*Endozoicomonas*-phage symbiosis.

At an MOI of 0.1, both phages showed an infection cycle of approximately 90 min, with EmPhiA exhibiting a longer latent period and a smaller burst size (60 min and 13.14 PFU/cell) than EmPhiS (30 min and 21.4 PFU/cell). These burst sizes are relatively small compared to those of other marine bacteriophages at even lower MOIs (0.01), such as phage HH109 (32.5 PFU/cell) in the marine pathogen *Vibrio alginolyticus* [44] and bacteriophage YC (500 PFU/cell) in the coral pathogen *V. coralliilyticus* [45]. This suggests that infection by EmPhiA and EmPhiS may not rapidly collapse the host bacterial population, which may allow continuous coexistence of both parties in the same host coral, making the phages potential core members of the holobiont. How corals regulate phage-bacterial population dynamics is essential in their symbiotic relationship and warrants further investigation.

Under TEM and cryo-EM observations, both EmPhiA and EmPhiS showed an icosahedral capsid and contractile tail, a morphology of the traditional *Myoviridae* morphotype A1 [46]. Notably, cryo-EM images revealed whiskers around the collar of each phage, which were invisible under TEM observation, highlighting the importance of cryo-EM in examining phage collar-whisker structure. The head size (∼130×122 nm) and head-to-tail length (∼390 nm) of EmPhiA and EmPhiS fall within the range of known jumbophages [47, 48]. Despite genome sizes smaller than the classical definition of jumbophages (>200 kb) [49], EmPhiA and EmPhiS have functionally related genes scattered in the genomes instead of modularly organized, consistent with other jumbophages such as LJR1-37 and φKZ [50, 51]. Both EmPhiA and EmPhiS also carry an NAD^+^ biosynthesis gene (NAMPT) and multiple tRNA genes (24 in EmPhiA, 25 in EmPhiS), another genomic feature common to jumbophages [47, 52]. Carrying their own genetic blueprint for NAD+ synthesis equips phages stronger resistance to the bacterial Thoeris defense system [53]. In addition, carrying tRNAs corresponding to codons more frequently used in their own genomes may redirect host cellular machinery to favor phage protein translation over host proteins [54, 55]. Supporting this notion, EmPhiA and EmPhiS possess more tRNA copies for serine, arginine, leucine, and methionine over others and employ some anticodons less frequent in the host tRNA gene pool. EmPhiA and EmPhiS thus constitute the first isolations of jumbophages that target non-pathogenic bacteria. Whether these phages are mutualistic or parasitic members of coral holobionts remains to be seen.

When compared with other prokaryotic dsDNA viruses, EmPhiA and EmPhiS are genetically distinct from other known viruses (1% coverage and ∼70% genome identity to the closest virus). The International Committee on Taxonomy of Viruses (ICTV) classifies viruses of the same species as having >95% nucleotide similarity. EmPhiA and EmPhiS (99.21% genome similarity) thus represent two strains of a novel viral species, which we name *Encorevirus taiwanensis*. Furthermore, phylogenetic analysis showed that EmPhiA and EmPhiS form an early-branching monophyletic group that separates from the neighboring *Demerecviridae* at an S_G_ value less than 0.005. Given that bacteriophage DSS3_VP1 (NC_055826) exhibited a similar S_G_ value in the proteomic tree and was proposed as a novel viral family, *Naomiviridae* [56], we propose that EmPhiA and EmPhiS also constitute a novel viral family, *Encoreviridae*, the ancient phylogenetic position of which may shed light on viral genomic diversity and evolution.

In addition to *E. montiporae* CL-33, both EmPhiA and EmPhiS infected *E. euniceicola* EF212. This is surprising as *E. montiporae* CL-33 was isolated from a stony coral, whereas *E. euniceicola* EF212 was from a soft coral [36, 37]. This suggests that EmPhiA and EmPhiS may be used by phylogenetically distant marine organisms to modulate their bacterial symbionts. A recent *Endozoicomonas* pan-genomic analysis showed that *E. montiporae* CL-33 and *E. euniceicola* EF212 form a monophyletic group [14]. Nevertheless, *E. gorgoniicola* PS125, which clusters with *E. montiporae* CL-33 and *E. euniceicola* EF212, was not infected by either phage. Whether the host specificity of EmPhiA and EmPhiS reflects coevolution with their bacterial hosts cannot be ascertained. Notably, despite their aforementioned similarities, chloroform sensitivity, temperature sensitivity, and one-step growth tests suggest that EmPhiA and EmPhiS have distinct physiological characters, such as envelope structure and life strategies. The phage envelope refers to a lipid layer derived from host cell membranes that surrounds the phage capsid and mediates penetration of phages into Gram-negative host bacteria [57, 58]. Thus, EmPhiA and EmPhiS probably utilize different infection mechanisms and may have distinct infection dynamics, even in the same host bacteria.

Using EmPhiS as an example, we examined host-phage transcriptomic dynamics underlying EmPhiS’s infection of *E. montiporae* CL-33. During the infection, the transcriptome slowly shifted from host to phage RNA, suggesting that EmPhiS gradually took over host transcription machinery. The early infection stage was marked by the highest phage gene expression and host DEGs. During this initial stage, phage EmPhiS expressed several genes to modulate host defense systems, e.g., antirepressors, cAMP PDE, and NAMPT. Phage antirepressors help prophage induction [59, 60]. Considering that *E. montiporae* CL-33 has 8 prophages in its genome [13], these antirepressors may constitute an EmPhiS countermeasure against phage superinfection exclusion. In addition, expression of the NAMPT gene allows EmPhiS to evade the bacterial Thoeris defense system, while expression of cAMP PDE may disrupt cyclic nucleotide homeostasis in the host bacterial cell [61], undermining bacterial defense system. Responding to the infection, host bacterial cells switch their energy metabolism from the TCA cycle to glycolysis and gluconeogenesis, together with downregulation of some non-vital functions, such as flagellar assembly. Bacterial PTS was also upregulated, likely enabling more efficient import of materials for glycolysis. A switch from aerobic to anaerobic respiration and employment of the glyoxylate shunt has been reported in several phage-infected bacteria [62–64]. This energy flow redirection may represent a general bacterial response to phage infection. Nevertheless, we found expression of phage MoeB at early infection stage and a Grx family protein at the subsequent middle stage. MoeB transfers sulfur for biosynthesis of molybdenum cofactor, which is vital for redox enzymes involved in diverse reactions [65, 66], while Grx protein catalyzes reversible oxidation/reduction of glutathione, an antioxidant tripeptide crucial for cellular redox homeostasis [67]. This phage gene expression is likely linked to upregulation of glutathione and cysteine metabolism in host cells. Cysteine is the precursor of glutathione. Together with this modulation of cellular glutathione homeostasis, the switch to anaerobic respiration in the host cell may alter host cellular oxidative status. Therefore, bacterial energy flow redirection during phage infection may also be a phage-driven response to create an anti-oxidative intracellular environment to promote phage infectivity [68, 69].

The middle infection stage corresponds to commencement of the phage lytic period and is highlighted by expression of phage DNA primase (DnaG) and several DNA-binding proteins. DNA primase is essential for initiation of DNA replication, whereas phage DNA-binding proteins may stabilize phage DNA during replication or may interrupt host DNA replication/repair [70, 71]. Together with preloading of other phage DNA replication genes (PolA, ligase, and nucleotidyltransferase) during the early stage, the middle infection stage likely marks the onset of phage genome replication. Notably, all middle stage genes were also expressed during the late stage (60 min), suggesting that the process of phage genome replication spans both the middle and late infection stages. In the host bacterium, in addition to continuous modulation of energy machinery, metabolisms of fatty acids many amino acids were suppressed. Reprogramming of host amino acid metabolism has been reported in phage-infected *Pseudomonas aeruginosa* and has been proposed as a strategy to prevent material depletion for phage replication [72, 73]. Accordingly, the middle infection stage may serve as a preparatory phase for subsequent synthesis of phage structural proteins.

The late infection stage assembles phage particles, as evidenced by TEM observations of phage particles in the host cell and expression of most phage morphogenesis genes at this stage. Concomitantly, two phage RNA polymerases, Rha, and homologs of bacterial Rho and RNA polymerase sigma-G factor, were expressed. The Rho factor contributes to transcription termination in bacterial cells [74], whereas phage-encoded sigma factors facilitate recognition of phage genes by bacterial RNA polymerases [75]. Expression of these auxiliary metabolic genes suggests that EmPhiS may promote transcription of its structure/morphogenesis genes both by expressing extra RNA polymerases and by redirecting host core transcriptional machinery. Corresponding to the final bursting of the host bacterial cell, at this stage we also observed expression of phage CwlK (a peptidoglycan LD-endopeptidase) and a TMhelix-containing protein, which degrade bacterial cell walls and disrupt cell membrane structure [76, 77]. In the host cell, cessation of most metabolic modulation in the early and middle stages was accompanied by upregulation of oxidative phosphorylation. Considering the reducing intracellular environment established during the early-middle stage, these late responses may generate oxidative stress, further destabilizing cell membranes to facilitate lysis of the host cell.

Notably, EmPhiS carries four tellurium-resistance protein-encoding genes (TerD, TerB, and TerZ), all of which were expressed during the early infection stage. Beside resistance to tellurite, diverse cellular functions have been proposed for tellurium-resistance proteins. For example, TerD domain-containing proteins possess Ca^2+^-binding activity and may be involved in responses to antibiotics, phage infection, and other external stressors [78]. In *Deinococcus radiodurans*, a multi-stressor-resistant bacterium, TerB is linked to resistance to stressors such as UVC irradiation and hydrogen peroxide [79, 80]. Disruption of TerB expression also reduces resistance of *Escherichia coli* K-12 to T5 phage infection [81].

Expression of these early genes may constitute a strategy for EmPhiS to establish its infection, which may provide the host bacterium stronger resistance to external stressors or other phages. To maximize reproductivity, many phages have evolved auxiliary metabolic genes to stabilize their hosts during the early infection stage, such as the psbA and psbD genes in cyanophages [82, 83]. However, to our knowledge, this is the first finding that phages may employ Ter family proteins as their infection weaponry. Intriguingly, the stress resistance conferred from EmPhiS infection may also stabilize the symbiosis between *Endozoicomonas* and corals under certain environmental conditions, rendering this phage infection a beneficial event for coral holobionts. Based on these findings, we propose an interaction model for EmPhiS infection of *E. montiporae* CL-33 and its putative contribution to coral holobionts (Fig. 8). Findings from this study offer new insights into complicated symbiotic interactions among phages, bacteria, and coral animals.

**Figure 8.**
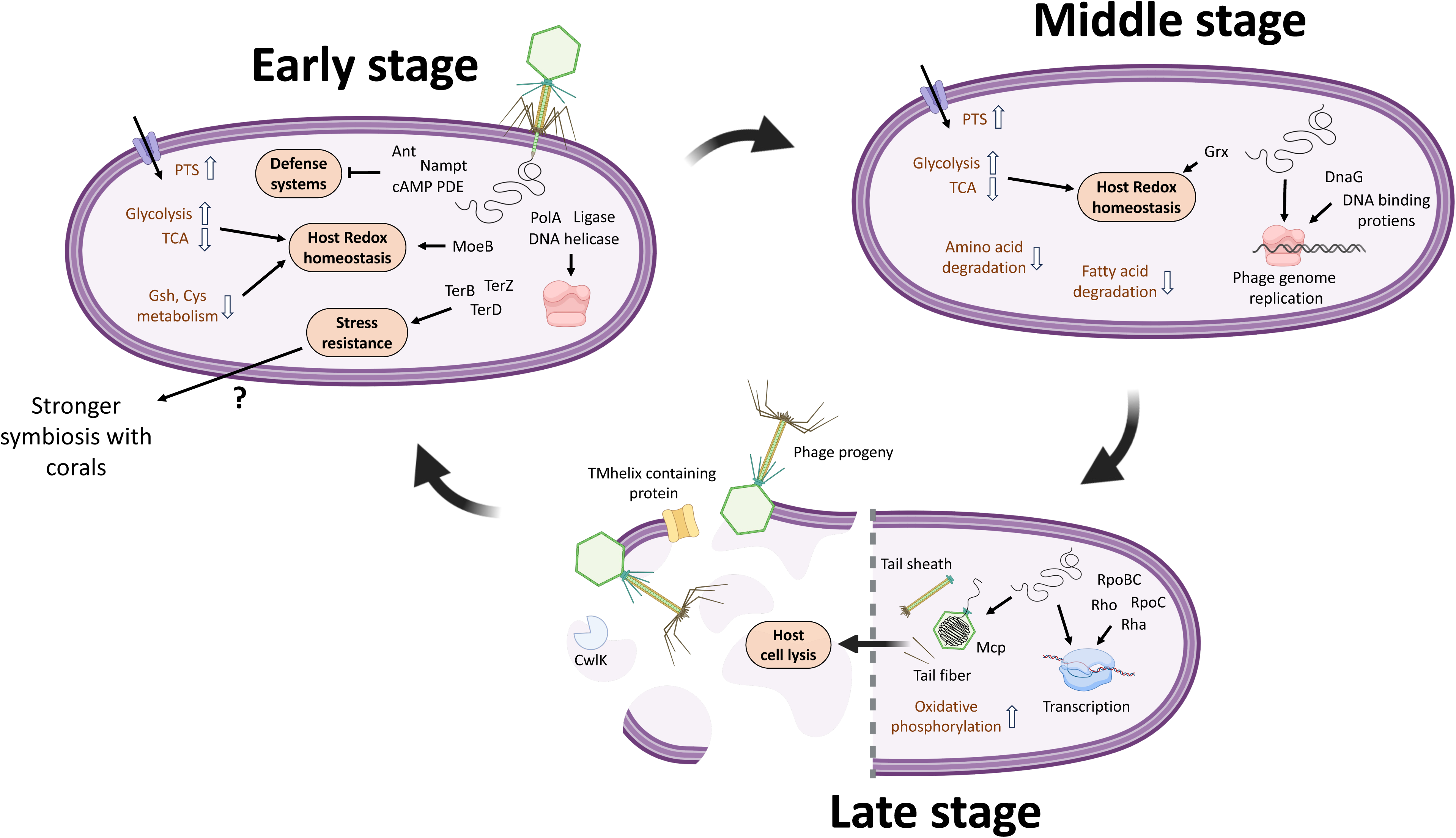
A model of phage-bacterium interaction at different stages of the infection cycle and putative contribution to coral holobionts. The early infection stage is highlighted by suppression of host defense systems, energy flow redirection, and modulation of intracellular redox status. All four Ter family genes are expressed at this stage, likely promoting host cell resistance to environmental stresses. The middle stage marks the onset of phage genome replication. The late stage is highlighted by expression of most phage structure proteins and lysis proteins, suggesting assembly of phage particles and bursting of the host cell. The slow propagation rate of EmPhiS may prevent collapse of host *Endozoicomonas* populations and may increase coral fitness in response to fluctuating environments.

## Acknowledgments

This research was funded by grants from the Academia Sinica, Taiwan (AS-IV-114-L03 and AS-PD-1141-L09-2). We thank Dr. Steven D. Aird for editing and commenting on this manuscript.

## Data availability

Data generated in this study are available on NCBI database under BioProject PRJNA1302128 and BankIt IDs 2996713 (EmPhiA genome assembly) and 2997144 (EmPhiS genome assembly).

---

**Figure S1.**
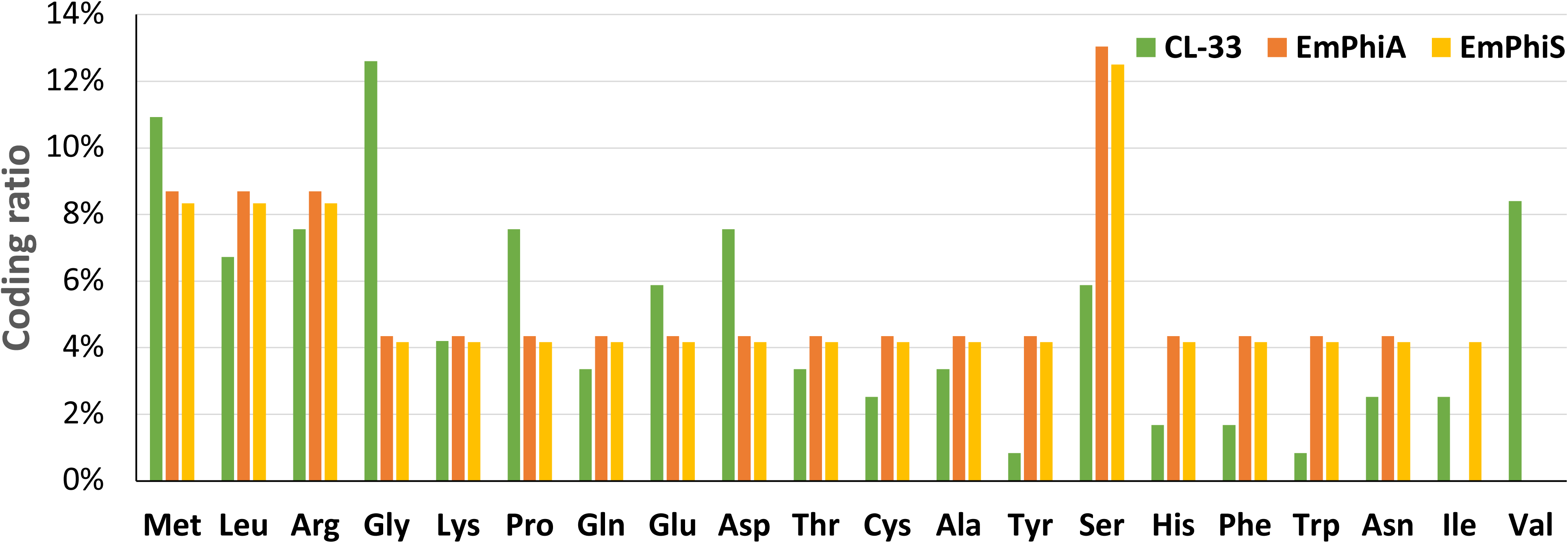
Relative coding ratios of 20 standard amino acids in EmPhiA, EmPhiS, and *Endozoicomonas montiporae* CL-33.

**Figure S2.**
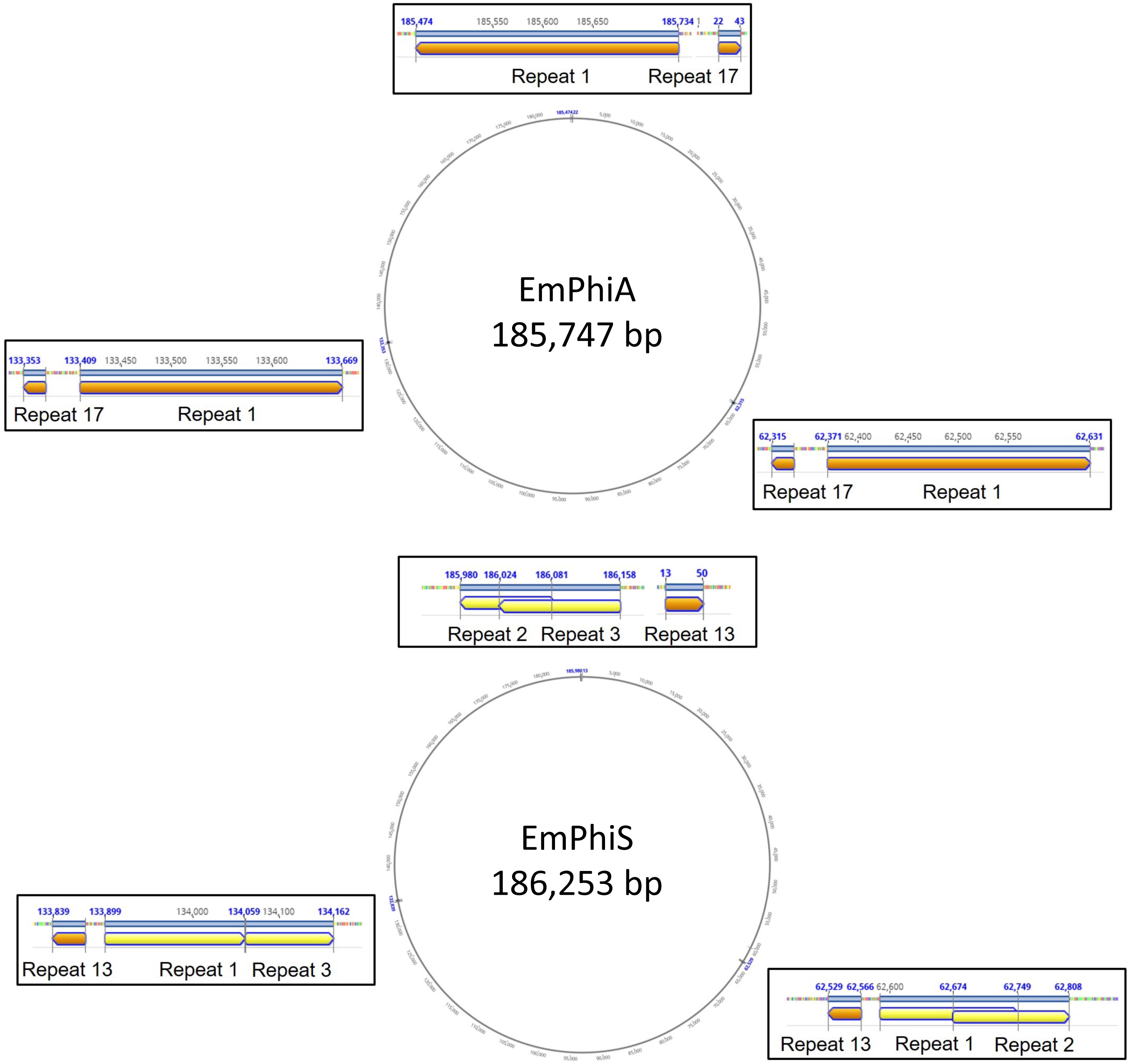
Long, repeated elements separate EmPhiA (top) and EmPhiS (bottom) genomes into three segments of approximately equal length.

**Figure S3.**
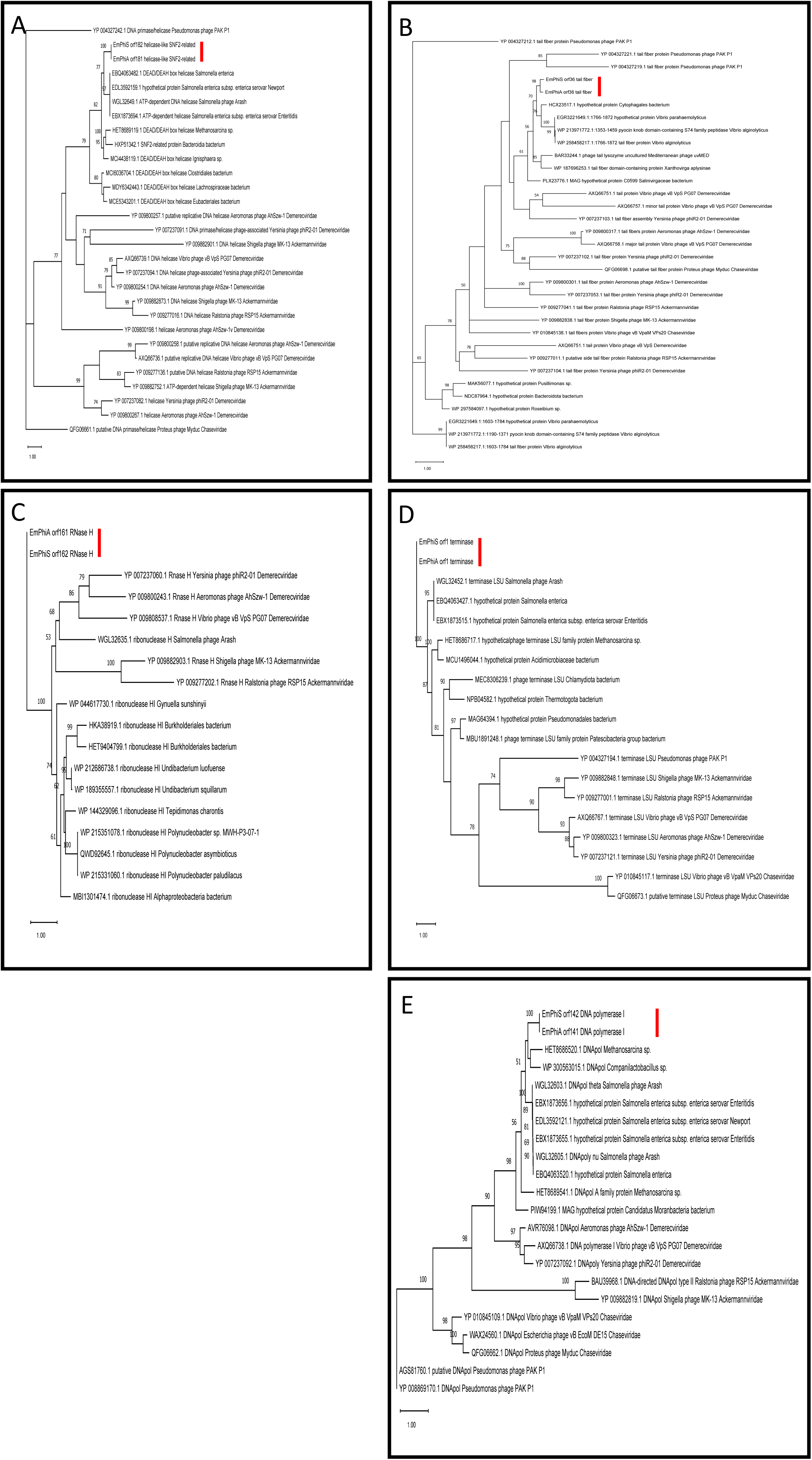
Marker gene-based phylogenetic analysis of EmPhiA and EmPhiS with *Pseudomonas* phage PAK_P1 and reference viruses from the viral families *Ackermannviridae*, *Chaseviridae*, and *Demerecviridae*. (a) helicase, (b) tail fiber, (c) RNase H, (d) terminase, (e) DNA polymerase. The monophyletic branch of EmPhiA and EmPhiS in each tree is highlighted with a red bar.

**Figure S4.**
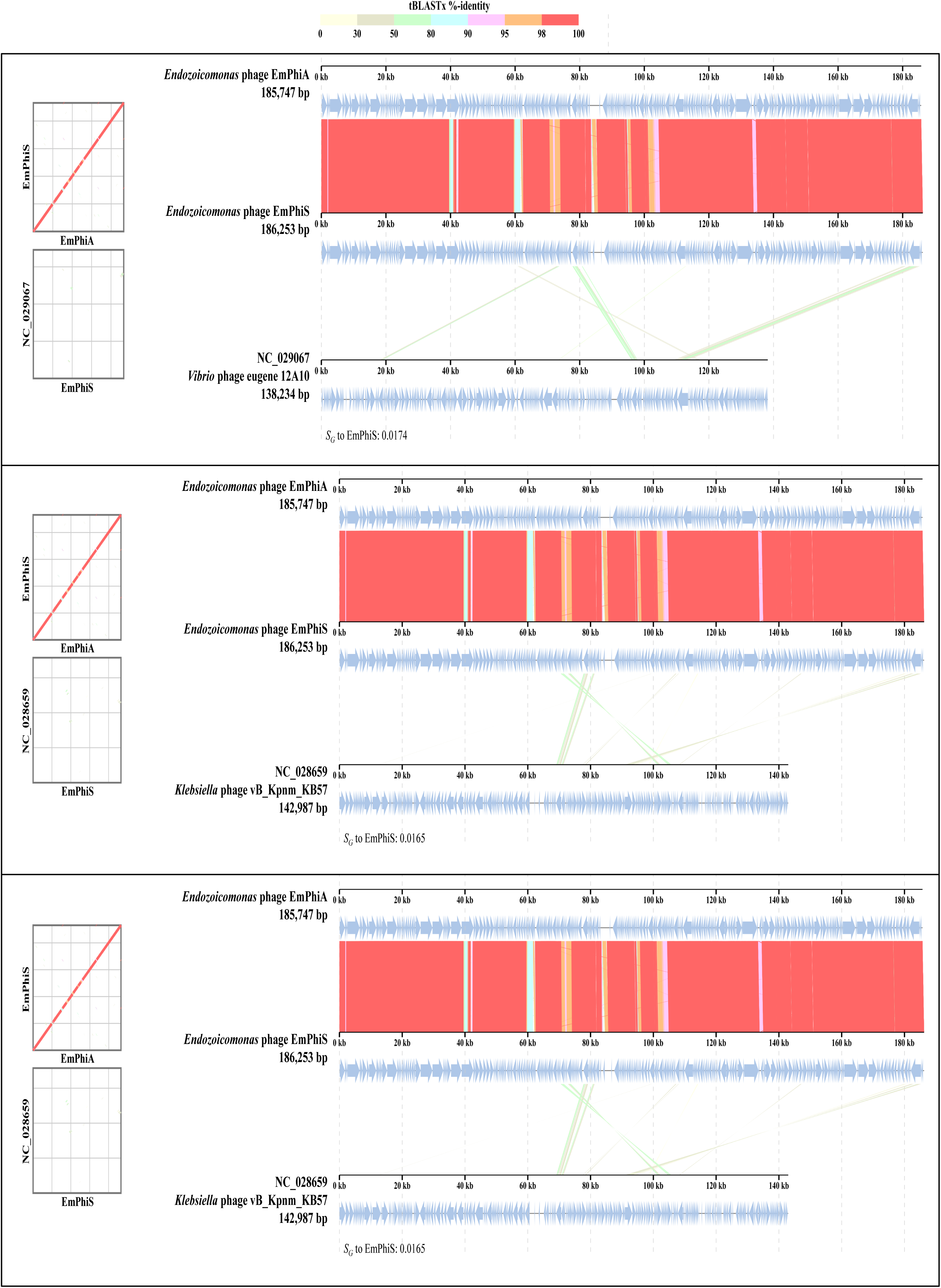
Genome alignment of EmPhiA and EmPhiS with the 3 most similar genomes among reference genomes in VIPtree.

**Figure S5.**
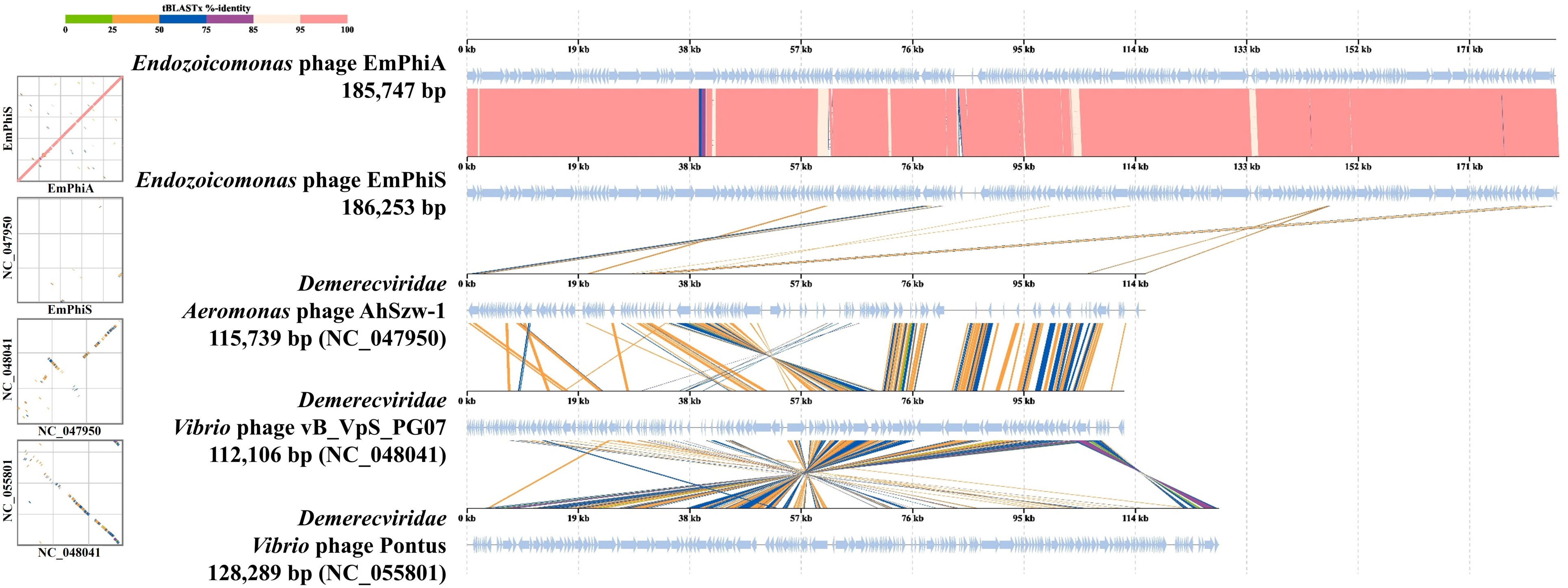
Genome alignment of EmPhiA and EmPhiS with 3 viruses in the *Demerecviridae*.

**Figure S6.**
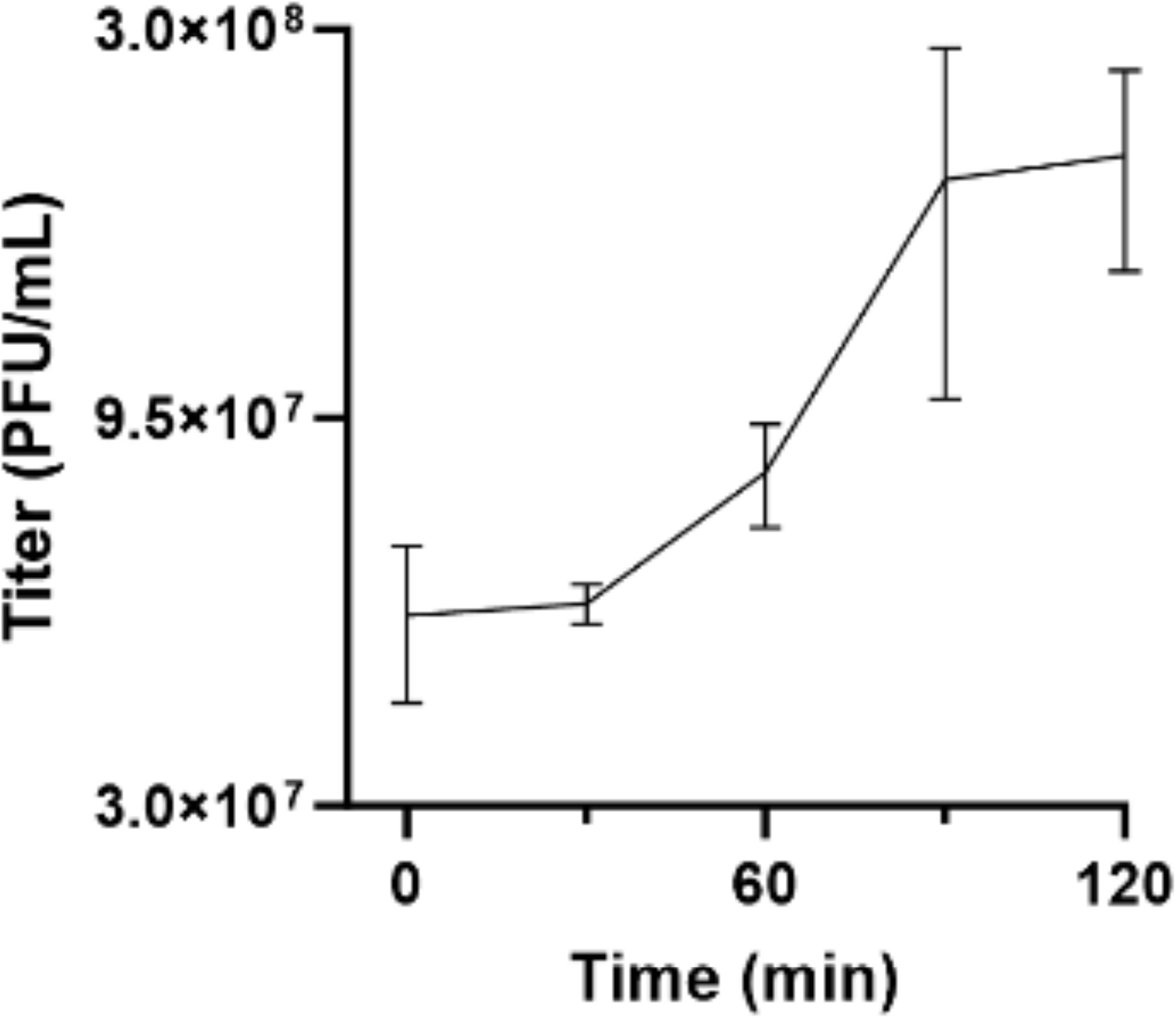
One-step growth curve of EmPhiS at a MOI of 3. Data are presented as averages and standard deviations of 5 replicates.

**Figure S7.**
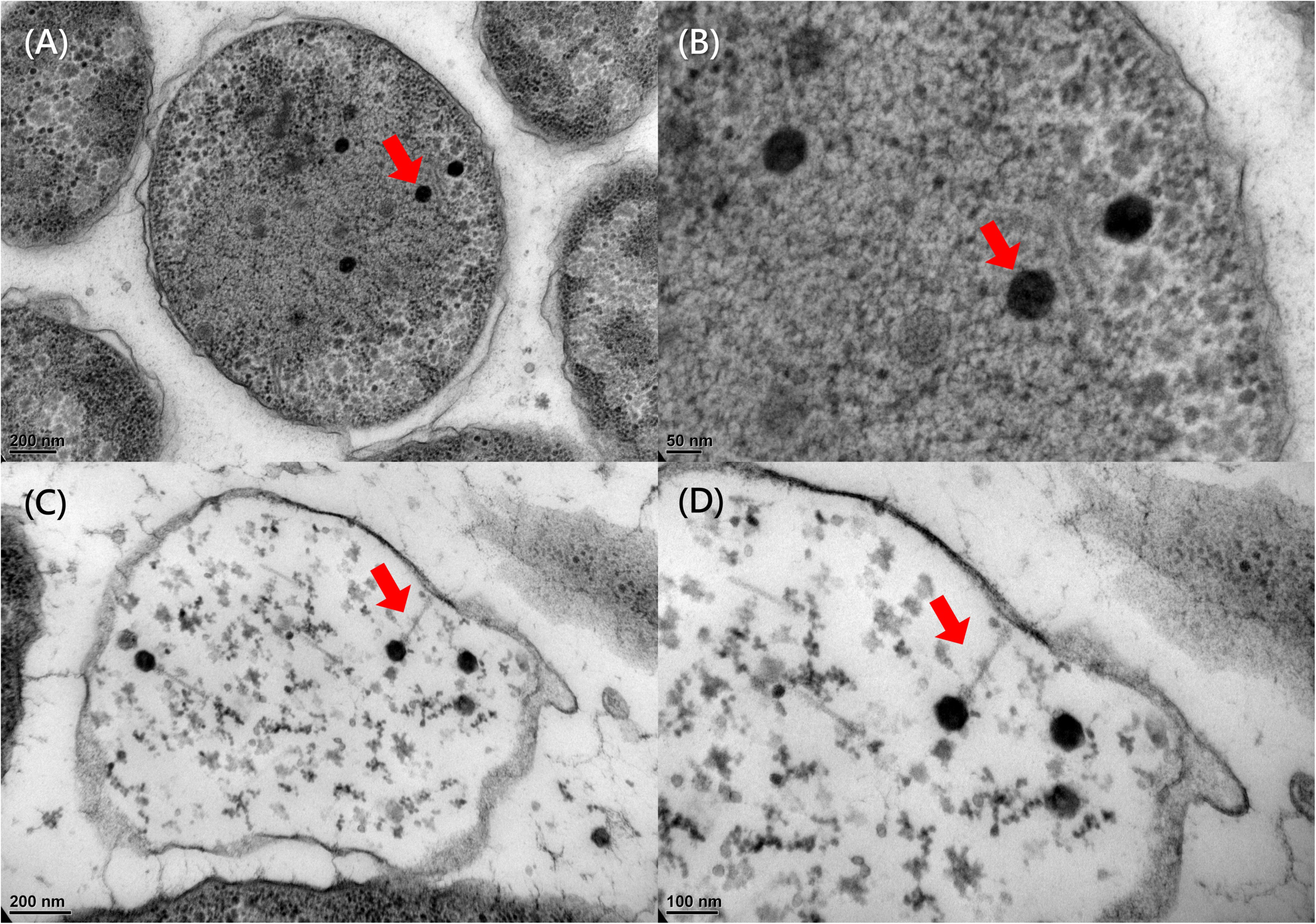
TEM images of an EmPhiS-infected bacterial cell. Red arrows highlight phage head structure (a,b) and assembled phage particles (c,d). (b) Close-up view of (a). (d) Close-up view of (c).

**Figure S8.**
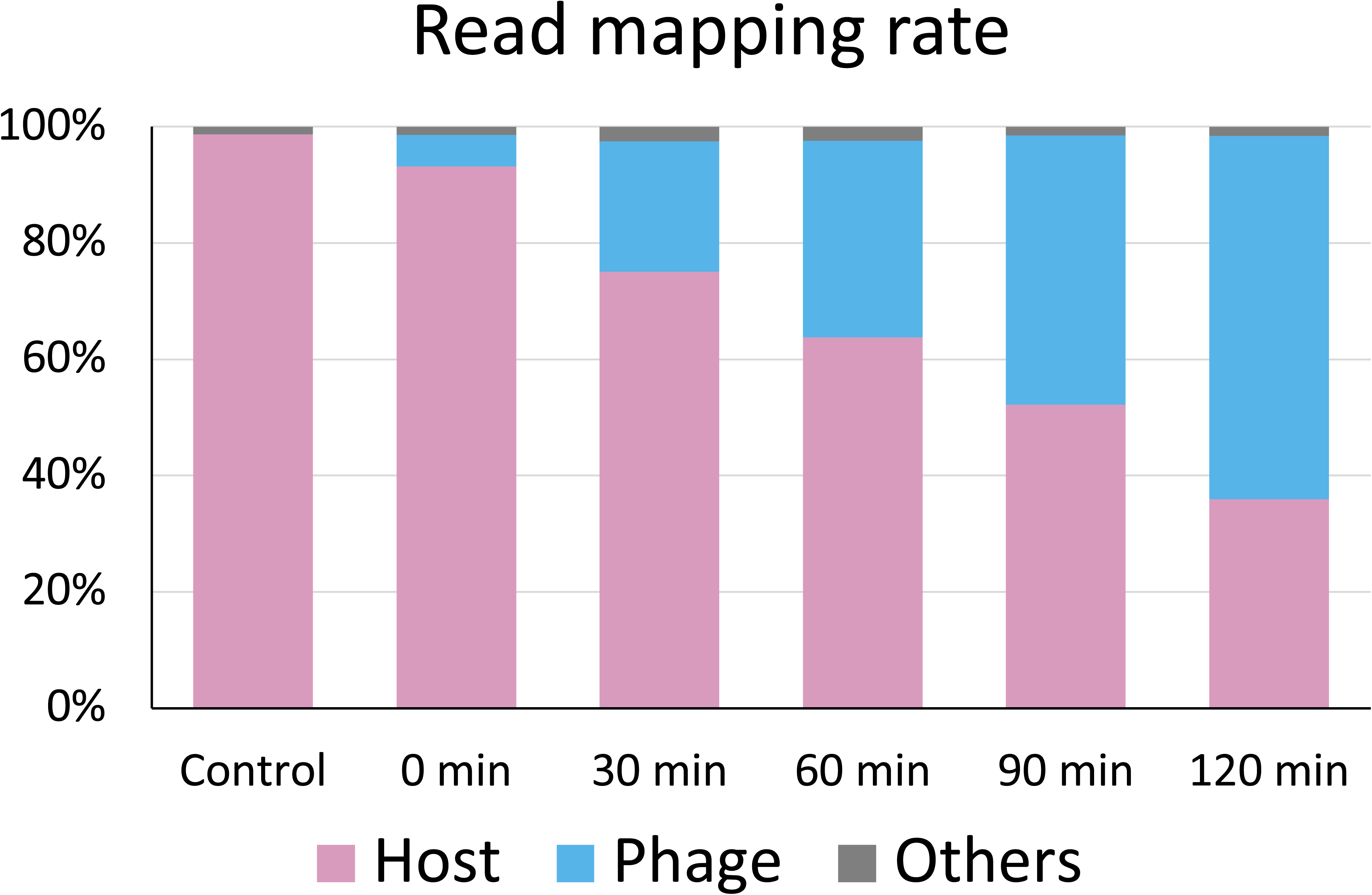
Relative ratios of phage and host reads in transcriptomic data during the infection experiment.

