## Supplementary Material 1 for "Bacteriophages of the predominant coral symbiont *Endozoicomonas*: novel models for coral holobiont interactions"

**Protocol of TEM sample cryo-fixation**

Phage-bacterial culture was centrifuged at 3,000 rpm at room temperature to separate infected bacteria and free phage particles. Bacterial pellet was placed into a 400 μm carrier and subjected to high-pressure freezing using a Leica EM PACT2 system at 2,000–2,050 bar. Frozen samples were then transferred to a Leica AFS2 system for freeze substitution, with a substitution solution comprising 1% OsO₄ and 0.1% uranyl acetate in acetone and a temperature gradient as follows:

| Temperature | Substitution period | temperature gradient |
| --- | --- | --- |
| −90 °C | 12 hours | +1°C/hour |
| −80 °C | 12 hours | +1°C/hour |
| −70 °C | 12 hours | +1°C/hour |
| −60 °C | 12 hours | +2°C/hour |
| −20 °C | 12 hours | +5°C/hour |
| 0 °C | 12 hours | +5°C/hour until 20°C |

At room temperature, samples were washed with acetone 2–3 times, each for 2 h. Infiltration was performed using Spurr’s resin (without DMAE/acetone) with a concentration gradient of 20%, 33%, 50%, 66%, 80%, 100%, 100%. Each step included microwaving at 250W for 2 min followed by 10-min shaking and another 2-min microwaving. Infiltrated samples were incubated at room temperature overnight, after which the resin was replaced with Spurr’s resin containing DMAE and incubated for 4 h. Embedded samples were polymerized at 70°C for 12–24 h and were sliced to 70-90 nm using a Leica Reichert Ultracut S or a Ultracut 7 ultramicrotome. TEM images were taken using an FEI Tecnai G2 Spirit Twin TEM. with a 4K x 4K Gatan Orius CCD camera, with a defocus range of -1.5 to -2.5 µm.
