## Supplementary Tables 1-5 for "Bacteriophages of the predominant coral symbiont *Endozoicomonas*: novel models for coral holobiont interactions"

**Supplementary Table 1.** ORFs with <90% identity between EmPhiA and EmPhiS.

| ORF | Location | | Length (bp) | Annotation | Identity  ( BLASTn) |
| --- | --- | --- | --- | --- | --- |
|  | Start | End |  |  |  |
| EM-phiA_36 | 38,925 | 42,764 | 3840 | Tail fiber | 84.56% |
| EM-phiS_36 | 38,925 | 42,701 | 3777 | Tail fiber |  |
| EM-phiA_73 | 62,066 | 62,296 | 231 | Hypothetical protein | EmPhiA unique |
| EM-phiS_70 | 61,582 | 62,103 | 522 | Phage regulatory protein | EmPhiS unique |
| EM-phiS_114 | 84,027 | 84,500 | 474 | HNH endonuclease |  |

**Supplementary Table 2.** Anticodon usage in phages EmPhiA, EmPhiS, and the host *Endozoicomonas montiporae* CL-33^T^ (NZ_CP013251.1). Numbers of anticodons in *E. montiporae* CL-33, EmPhiA, and EmPhiS are delimited by slashes. Grey boxes highlight anticodons employed by both phages and the host bacterium. Anticodons are predicted using tRNAScan-SE v.2.0.7. Anticodons for Supres (CTA, TTA, TAC) and SelCys (TCA) were not detected. N-formylmethionine (fMet) in *E. montiporae* CL-33 was counted into Met.

| **Number of the anticodons of the CL-33^T^/ phiA/ phiS** | | | | | | |
| --- | --- | --- | --- | --- | --- | --- |
| **Amino acid** | **Isotype of anticodon** | | | | | |
| **Ala: 4/1/1** | AGC: - | GGC: 3/0/0 | CGC: - | **TGC: 1/1/1** |  |  |
| **Gly: 15/1/1** | ACC: - | GCC: 14/0/0 | CCC: - | **TCC: 1/1/1** |  |  |
| **Pro: 9/1/1** | AGG: - | GGG: 2/0/0 | CGG: 1/0/0 | **TGG: 6/1/1** |  |  |
| **Thr: 4/1/1** | AGT: - | GGT: 1/0/0 | CGT: 2/0/0 | **TGT: 1/1/1** |  |  |
| **Val: 1/0/0** | AAC: - | GAC: 1/0/0 | CAC: - | TAC: 9/0/0 |  |  |
| **Ser: 7/3/3** | AGA: - | **GGA: 2/1/1** | CGA: - | **TGA: 3/1/1** | ACT: - | **GCT: 2/1/1** |
| **Arg: 9/2/2** | **ACG: 6/1/1** | GCG: - | CCG: 1/0/0 | TCG: - | CCT: 1/0/0 | **TCT: 1/1/1** |
| **Leu: 8/2/2** | AAG: - | GAG: 1/0/0 | CAG: 2/0/0 | **TAG: 1/1/1** | CAA: 2/0/0 | **TAA: 2/1/1** |
| **Phe: 2/1/1** | AAA: - | **GAA: 2/1/1** |  |  |  |  |
| **Asn: 3/1/1** | ATT: - | **GTT: 3/1/1** |  |  |  |  |
| **Lys: 5/1/1** |  |  | CTT: - | **TTT: 5/1/1** |  |  |
| **Asp: 9/1/1** | ATC: - | **GTC: 9/1/1** |  |  |  |  |
| **Glu: 7/1/1** |  |  | CTC: - | **TTC: 7/1/1** |  |  |
| **His: 2/1/1** | ATG: - | **GTG: 2/1/1** |  |  |  |  |
| **Gln: 4/1/1** |  |  | CTG: - | **TTG: 4/1/1** |  |  |
| **Ile: 3/0/1** | AAT: - | GAT: 1/0/1 | CAT: 2/0/0 | TAT: - |  |  |
| **Met: 13/2/2** |  |  | **CAT: 13/2/2** |  |  |  |
| **Tyr: 1/1/1** | ATA: - | **GTA: 1/1/1** |  |  |  |  |
| **Cys: 3/1/1** | ACA: - | **GCA: 3/1/1** |  |  |  |  |
| **Trp: 1/1/1** |  |  | **CCA: 1/1/1** |  |  |  |

**Supplementary Table 3.** EmPhiS early infection genes.

| Functional category | Gene ID & Gene product |
| --- | --- |
| **Infection and lysis** | EmPhiS_45: Antirepressor  EmPhiS_59: YrbG  EmPhiS_132: Hydrolase  EmPhiS_138: Antirepressor N-terminal domain-containing protein  EmPhiS_140: Antirepressor N-terminal domain-containing protein  EmPhiS_186: 3',5'-cyclic AMP phosphodiesterase protein |
| **Nucleotide metabolism, recombination, replication** | EmPhiS_26: KilA-N domain-containing protein  EmPhiS_49: Macro domain-containing protein  EmPhiS_60: TerB  EmPhiS_61: TerZ,  EmPhiS_63: TerD  EmPhiS_64: TerD  EmPhiS_82: Polynucleotide kinase  EmPhiS_86: RNA ligase 1  EmPhiS_91: Nucleotidyltransferase  EmPhiS_111: Ribose-phosphate pyrophosphokinase  EmPhiS_142: PolA  EmPhiS_152: RNA ligase family protein  EmPhiS_156: DNA ligase  EmPhiS_159: KilA-N domain-containing protein  EmPhiS_160: 3' exoribonuclease  EmPhiS_162: RnaseH1  EmPhiS_166: ATP-dependent DNA helicase  EmPhiS_187: KilA-N domain-containing protein  EmPhiS_193: KilA-N domain-containing protein  EmPhiS_240: KilA-N domain-containing protein  EmPhiS_246: Deoxynucleotide monophosphate kinase  EmPhiS_249: Ribonucleoside-diphosphate reductase 1 subunit beta |
| **Energy metabolism** | EmPhiS_106: Nampt  EmPhiS_118: MFS transporter  EmPhiS_119: UroD  EmPhiS_121: Coil containing protein  EmPhiS_163: HAD domain-containing protein  EmPhiS_238: MoeB |
| **Structure assembly & morphogenesis** | EmPhiS_105: BCS1 |

**Supplementary Table 4.** EmPhiS middle infection genes.

| Functional category | Gene ID & gene product |
| --- | --- |
| **Infection and lysis** | EmPhiS_9: Cell wall hydrolase |
| **Nucleotide metabolism, recombination, replication** | EmPhiS_137: HupB  EmPhiS_161: DnaG  EmPhiS_169: ssDNA binding protein  EmPhiS_174: RpoD  EmPhiS_182: ATP-dependent DNA helicase  EmPhiS_220: DNA-binding domain-containing protein  EmPhiS_250: Deoxynucleoside kinase  EmPhiS_253: Ribonucleoside-diphosphate reductase 1 subunit alpha  EmPhiS_254: Grx family protein |
| **Energy metabolism** | EmPhiS_39: Twitch domain-containing radical SAM aprotein  EmPhiS_40: Twitch domain-containing radical SAM protein  EmPhiS_41: Twitch domain-containing radical SAM protein  EmPhiS_42: Twitch domain-containing radical SAM protein |
| **Structure assembly & morphogenesis** | EmPhiS_36: Tail fiber protein  EmPhiS_181: Porin |

**Supplementary Table 5.** EmPhiS late infection genes.

| Functional category | Gene ID & gene product |
| --- | --- |
| **Infection and lysis** | EmPhiS_18: CwlK  EmPhiS_29: Peptidase S74 domain-containing protein  EmPhiS_231: Putative internalin  EmPhiS_234: SEA domain containing protein  EmPhiS_235: TMhelix containing protein |
| **Nucleotide metabolism, recombination, replication** | EmPhiS_11: RNA polymerase sigma-G factor  EmPhiS_69: Thymidine kinase  EmPhiS_71: Rha  EmPhiS_72: Repair protein  EmPhiS_116: HNH endonuclease  EmPhiS_191: RpoBC  EmPhiS_196: RpoC  EmPhiS_207: Endonuclease  EmPhiS_208: HNH endonuclease  EmPhiS_210: Rho |
| **Energy metabolism** | EmPhiS_30: LamG-like jellyroll fold domain-containing protein  EmPhiS_74: GlgX |
| **Structure assembly & morphogenesis** | EmPhiS_1: Terminase, large subunit  EmPhiS_28: Tail fiber protein  EmPhiS_115: Tail fiber  EmPhiS_189: Portal  EmPhiS_206: Mcp  EmPhiS_215: Tail sheath protein  EmPhiS_219: Baseplate puncturing device |
